# YAP and TAZ couple osteoblast precursor mobilization to angiogenesis and mechanoregulated bone development

**DOI:** 10.1101/2023.01.20.524918

**Authors:** Joseph M. Collins, Annemarie Lang, Cristian Parisi, Yasaman Moharrer, Madhura P. Nijsure, Jong Hyun (Thomas) Kim, Greg L. Szeto, Ling Qin, Riccardo L. Gottardi, Nathanial A. Dyment, Niamh C. Nowlan, Joel D. Boerckel

## Abstract

Endochondral ossification requires coordinated mobilization of osteoblast precursors with blood vessels. During adult bone homeostasis, vessel adjacent osteoblast precursors respond to and are maintained by mechanical stimuli; however, the mechanisms by which these cells mobilize and respond to mechanical cues during embryonic development are unknown. Previously, we found that deletion of the mechanoresponsive transcriptional regulators, YAP and TAZ, from Osterix-expressing osteoblast precursors and their progeny caused perinatal lethality. Here, we show that embryonic YAP/TAZ signaling couples vessel-associated osteoblast precursor mobilization to angiogenesis in developing long bones. Osterix-conditional YAP/TAZ deletion impaired endochondral ossification in the primary ossification center but not intramembranous osteogenesis in the bone collar. Single-cell RNA sequencing revealed YAP/TAZ regulation of the angiogenic chemokine, Cxcl12, which was expressed uniquely in vessel-associated osteoblast precursors. YAP/TAZ signaling spatially coupled osteoblast precursors to blood vessels and regulated vascular morphogenesis and vessel barrier function. Further, YAP/TAZ signaling regulated vascular loop morphogenesis at the chondro-osseous junction to control hypertrophic growth plate remodeling. In human cells, mesenchymal stromal cell co-culture promoted 3D vascular network formation, which was impaired by stromal cell YAP/TAZ depletion, but rescued by recombinant CXCL12 treatment. Lastly, YAP and TAZ mediated mechanotransduction for load-induced osteogenesis in embryonic bone.

## Introduction

The appendicular skeleton forms by the conversion of avascular cartilage templates to vascularized bones via endochondral ossification. In long bones, endochondral ossification initiates in the primary ossification center upon coordinated invasion by osteoblast precursors and blood vessels. Osterix (Osx)-expressing osteoblast precursors originate in the surrounding intramembranous bone collar and co-invade with blood vessels as pericytes (*1*). Specialized bone blood vessels, composed of type H endothelial cells, mediate cartilage matrix remodeling and localize with vessel-associated osteoblast precursors to support new bone formation (*2, 3*). Upon invasion, these precursors either remain vessel-associated to support angiogenesis or mature into osteoblasts for bone formation. During adult bone homeostasis, vessel adjacent osteo-lineage precursors respond to and are maintained by mechanical stimuli (*4*); however, the mechanisms by which these cells mobilize and respond to mechanical cues during embryonic development are unknown. Here, we show that the mechanoresponsive transcriptional regulators, Yes-associated protein (YAP) and transcriptional co-activator with PDZ-binding motif (TAZ), couple osteoblast precursors to vessel morphogenesis and mediate mechanical regulation of fetal bone development.

During human fetal development, insufficient fetal movement (akinesia) causes skeletal defects. Fetal akinesia, caused by genetic defects in muscle development, insufficient amniotic fluid volume, or restricted movement, can lead to joint dysplasia (e.g., developmental dysplasia of the hip, arthrogryposis) and impaired fetal bone formation (*5*). Normal fetal movement places high stress and strain on regions associated with akinesia-impaired morphogenesis (*6*), suggesting that proper skeletal development requires movement-induced mechanical cues. How these mechanical cues are transduced by skeletal cells to direct morphogenesis is poorly understood. YAP and TAZ mediate mechanotransduction in osteoblast precursors *in vitro* (*7*), are required for persistent cell motility (*8*), and combinatorially regulate bone development (*9, 10*). Deletion of YAP and/or TAZ from Osx-expressing cells and their progeny causes perinatal lethality and severe bone fragility in mice (*9*). YAP and TAZ are activated by a variety of signals, including morphogenic and mechanical cues, and regulate gene expression by transcriptional co-activation/repression of other transcription factors (*11*). These data position YAP and TAZ as potential mediators of both embryonic bone morphogenesis and mechanotransduction.

Here, we show that YAP/TAZ signaling in osteoblast-lineage cells mediates bone development by regulating the co-mobilization of vessel-associated osteoblast precursors and blood vessels through CXCL12 signaling. YAP/TAZ-mediated vascular morphogenesis is necessary for hypertrophic cartilage degradation and endochondral ossification. Further, YAP and TAZ are required for embryonic load-induced bone formation.

## Results

### YAP and TAZ mediate endochondral ossification and growth plate remodeling

We first sought to determine the roles of YAP and TAZ in embryonic bone development. In the developing bone, YAP and TAZ are expressed robustly in cells of the primary ossification center, particularly the Osterix-expressing osteoblast precursors (Fig. 1A). We therefore deleted both YAP and TAZ from Osterix-expressing cells using Osx1-GFP::Cre. Osx is predominately expressed in hypertrophic chondrocytes and osteoprogenitors and their progeny during embryonic limb development (Fig. 1A). Cre-mediated recombination effectively deleted YAP and TAZ from Osx-expressing cells in both the primary ossification center and bone collar but did not affect YAP/TAZ expression in the cartilage rudiment, blood vessels, or surrounding muscle (Fig. 1A). To assess early osteoblast activity, we measured alkaline phosphatase (ALP) activity, which marks osteogenesis in both endochondral ossification in the primary ossification center and intramembranous ossification in the bone collar. Osx-conditional YAP/TAZ deletion significantly reduced ALP activity in the primary ossification center (Fig. 1B,C), but did not substantially alter ALP activity or mineralization in the bone collar (Fig. 1B,C, Fig. S1). These data suggest specific roles for YAP and TAZ in endochondral ossification.

**Figure 1.**
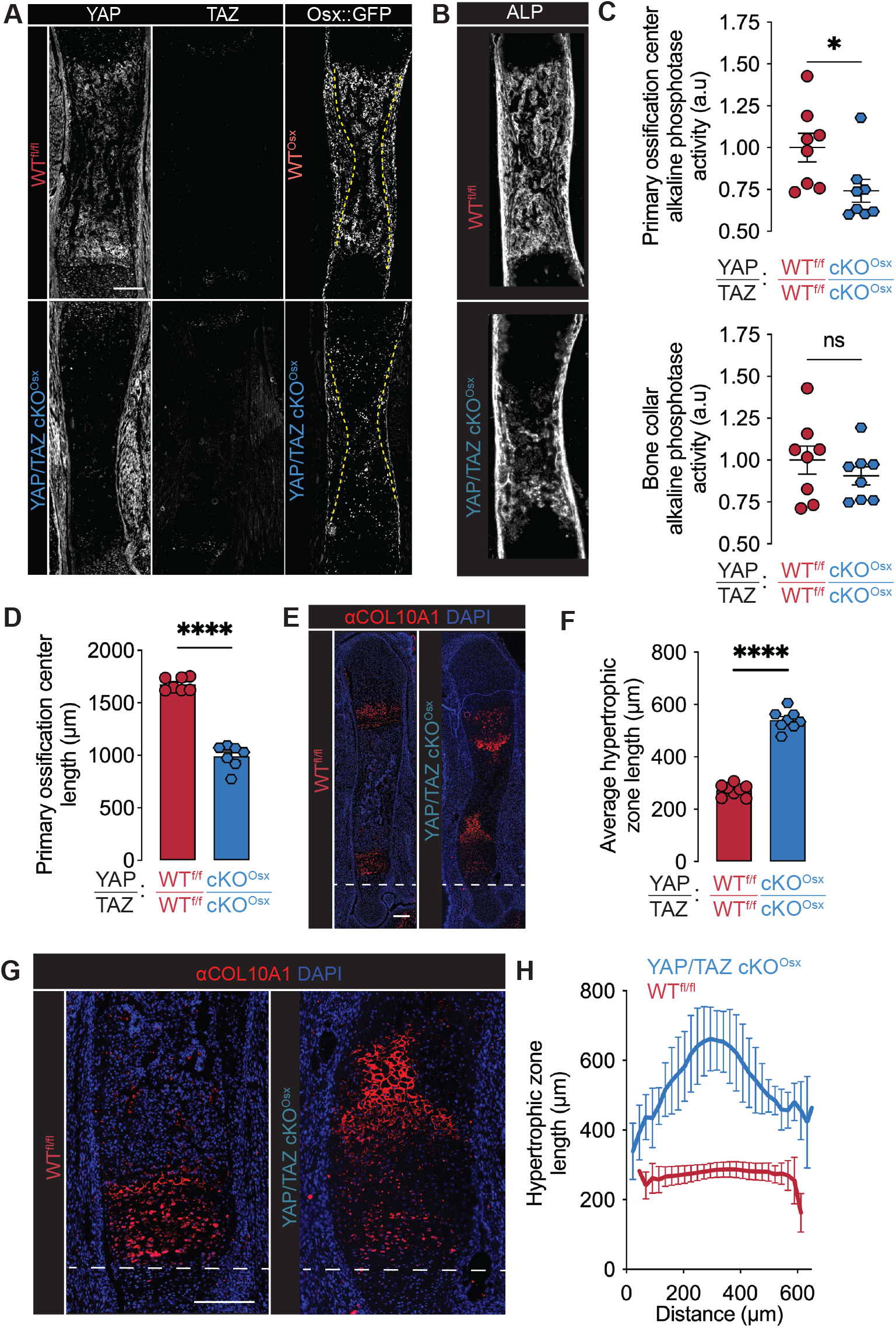
YAP/TAZ mediate endochondral ossification in fetal limb development. (**A**) YAP, TAZ, Osx::GFP immunostains in E17.5 humeri (n= 2-4). Dotted lines mark the boundary between primary ossification center and bone collar (**B**) Representative alkaline phosphatase (ALP) activity stain in E17.5 femurs (n=8). (**C**) Quantification of ALP activity stain by region of interest. (**D**) Quantification of primary ossification center length. (**E**) Collagen-10 stain in WT^fl/fl^ and YAP/TAZ cKO^Osx^ E17.5 femurs (n=8). (**F**) Quantification of average growth plate length. (**G**) Zoomed images of the fetal growth plate stained for Collagen-10. (**H**) Quantification of growth plate morphology (n = 8). Scale bars: 200 μm. 200 μm. Error bars: SEM, except SD in panel H. ‘*’: p<0.05; ‘****’: p<0.0001.

Endochondral ossification requires chondrocyte hypertrophy and hypertrophic cartilage remodeling. Upon hypertrophy, chondrocytes secrete a Collagen 10-rich matrix, which is degraded by endothelial cells and other cartilage-degrading cells to form the transverse cartilage septum at the chondro-osseous junction. In the primary spongiosa, the matrix is replaced by osteoblast-lineage cells. Osx-conditional YAP/TAZ deletion significantly decreased the length of the primary ossification center (Fig. 1D). Osterix-conditional YAP/TAZ deletion increased the length of the growth plate (Fig. 1E,F). YAP/TAZ deletion did not alter chondrocyte entry into hypertrophy, marked by the initiation of Collagen-10 expression, but expanded the length of the hypertrophic zone. Notably, YAP/TAZ deletion altered the shape of the cartilage septum, producing a cone-shaped chondro-osseous junction that extended hypertrophic cartilage into the primary ossification center (Fig. 1G,H). Together, these data indicate that YAP and TAZ mediate endochondral ossification and hypertrophic cartilage remodeling.

### YAP and TAZ mediate osteoblast precursor mobilization

Endochondral ossification requires osteoblast precursors to co-mobilize with blood vessels from the surrounding bone collar into the primary ossification center. Cells move by exerting forces on their extracellular matrix through cytoskeletal activation (*8*). We mapped skeletal-lineage cells over time using triple collagen reporter mice, which express fluorophores that label chondrocytes (Col2-CFP), hypertrophic chondrocytes (Col10-RFP), and osteoblasts (Col1(3.6kb)-YFP) (Fig. 2A). Anlage chondrocytes reduced Col2-CFP expression upon maturation into Col10-RFP-expressing hypertrophic chondrocytes (Fig. 2B). Growth plate separation, indicated by distinct Col10-RFP peaks, initiated at E15.5 and continued through E17.5 (Fig. 2B). At E15.5, differentiated Col1(3.6kb)-YFP^+^ osteoblasts were present in the bone collar, and appeared in the primary ossification center by E17.5. To map the cytoskeleton, we stained filamentous actin (F-actin) using Phalloidin (Fig. 2B, Fig. S2). An acute peak in F-actin intensity emerged in the middle of the primary ossification center at E15.5, marking the initiation of cellular mobilization (Fig. 2B). By E17.5, elevated F-actin staining extended as a uniform plateau throughout the primary ossification center, coincident with Col1(3.6kb)-YFP^+^ osteoblast emergence (**Fig. 2B, Fig. S2**).

**Figure 2.**
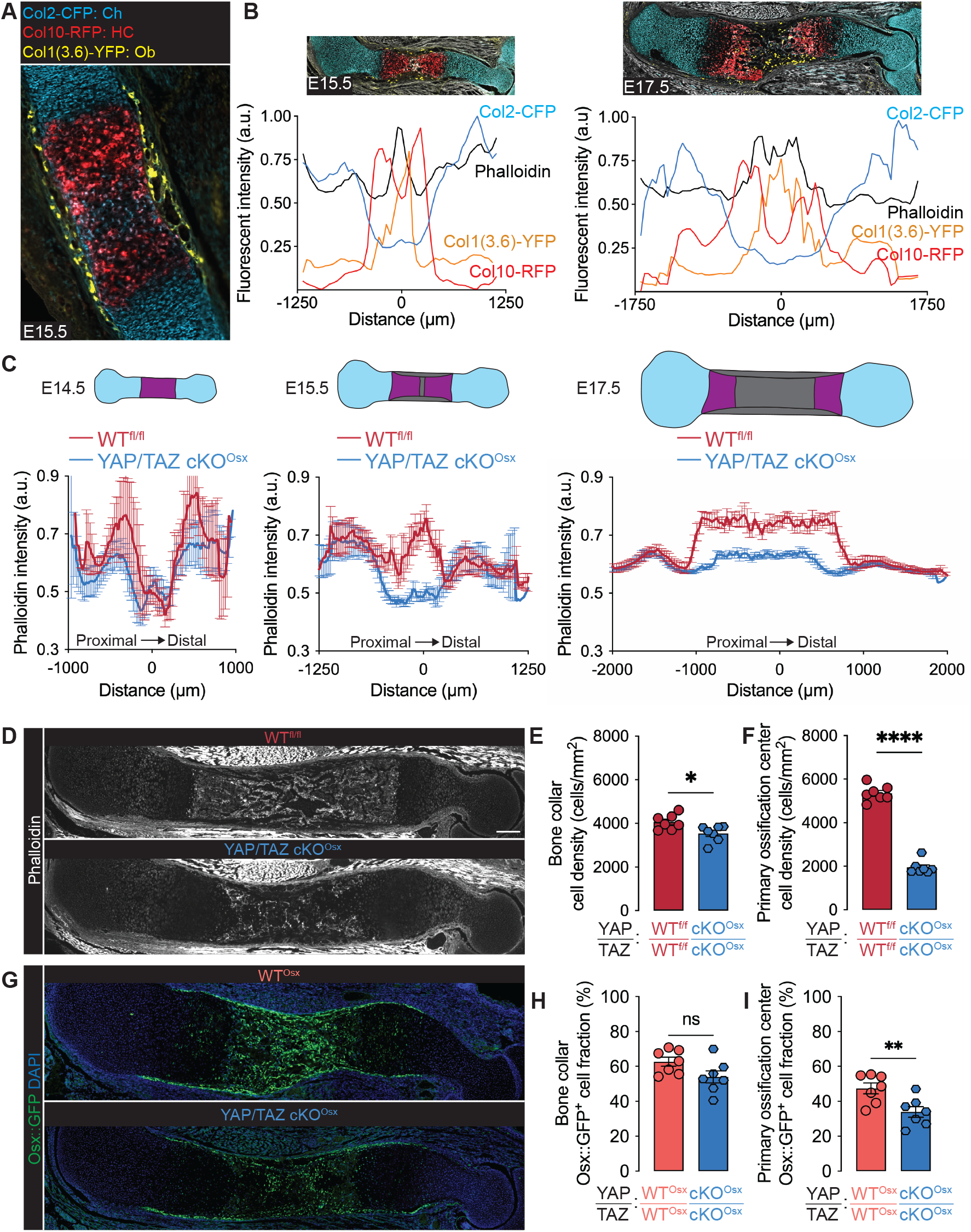
YAP/TAZ mediate osteoblast precursor localization and cytoskeletal dynamics in fetal bone development. (**A**) E15.5 tibia from Col2-CFP; Col10-RFP; Col1(3.6kb)-YFP transgenic reporter. (**B**) Relative fluorescent reporter and phalloidin intensity along primary axis of triple reporter humeri at E15.5 and E17.5 (n=3). Images scaled with x-axis. (**C**) Quantification of Phalloidin intensity along the longitudinal humeral axis of WT^fl/fl^ and YAP/TAZ cKO^Osx^ limbs (E14.5: n =3-4, E15.5: n=6, E17.5: n=7). Error bars: SEM. (**D**) Representative phalloidin image in E17.5 humeri. (**E**,**F**) Quantification of cells per area by DAPI in regions of interest in E17.5 femurs. (**G**) Representative image of Osx:GFP and DAPI in E17.5 humeri. (**H**,**I**) Fraction of Osx::GFP-expressing cells in regions of interest in 17.5 femurs. Scale bar: 200 μm. Error bars: SEM. ‘*’: p<0.05; ‘**’: p<0.01; ‘****’: p<0.0001.

YAP and TAZ are activated in migrating cells by tension in the actin cytoskeleton (*8*). Osx-conditional YAP/TAZ deletion had no effect on F-actin distribution at E14.5 (Fig. 2C, S3), but prevented the emergence of the mid-diaphyseal F-actin peak at E15.5, indicating a delay in primary ossification center initiation (Fig. 2C, Fig. S3). At E17.5, Osx-conditional YAP/TAZ deletion both shortened and reduced the F-actin plateau that marks primary ossification center expansion (Fig. 2C,D). Next, to directly evaluate osteoblast precursor emergence, we quantified cell density and the fraction of Osx::GFP^+^ cells in the bone collar and primary ossification center. Osx-conditional YAP/TAZ deletion substantially reduced total cell density in the primary ossification center, but only modestly reduced total cell density in the bone collar (Fig. 2E,F). YAP/TAZ deletion significantly reduced Osx::GFP^+^ cell fraction in the primary ossification center but did not significantly alter the fraction of Osx::GFP^+^ cells in the bone collar (Fig. 2G-I, Fig. S4). These data suggest that YAP and TAZ mediate osteoblast precursor mobilization into the developing limb.

### YAP and TAZ regulate Cxcl12 in vessel-associated osteoblast precursors

Vessel-associated osteoblast precursors (VOPs) mobilize with invading blood vessel, prior to their commitment to mature osteoblastic fate (*1*). Therefore, to determine how YAP and TAZ regulate osteoblast-lineage cell transcription and their interaction with other cell types, we performed single-cell RNA sequencing (scRNA-seq) on embryonic forelimbs. We isolated cells from E17.5 forelimbs of WT^fl/fl^, WT^Osx^, and YAP/TAZ cKO^Osx^ mice (n = 3-4/genotype) for scRNA-seq using the 10X Genomic Chromium platform. To capture both osteoblast-lineage and non-osteoblast-lineage cells, we removed dead cells and lysed red blood cells, but did not sort prior to scRNA-seq. We evaluated high-quality transcriptomes from 120,292 cells (Fig. 3A). The developing embryonic limb contains a large number of cell types. We applied Louvain clustering to find 77 clusters, categorized as 12 major cell populations by canonical marker expression (*12, 13*). These populations included osteoblasts, endothelial cells, and chondrocytes (ss**Fig. 3B, Fig. S5-11**).

**Figure 3.**
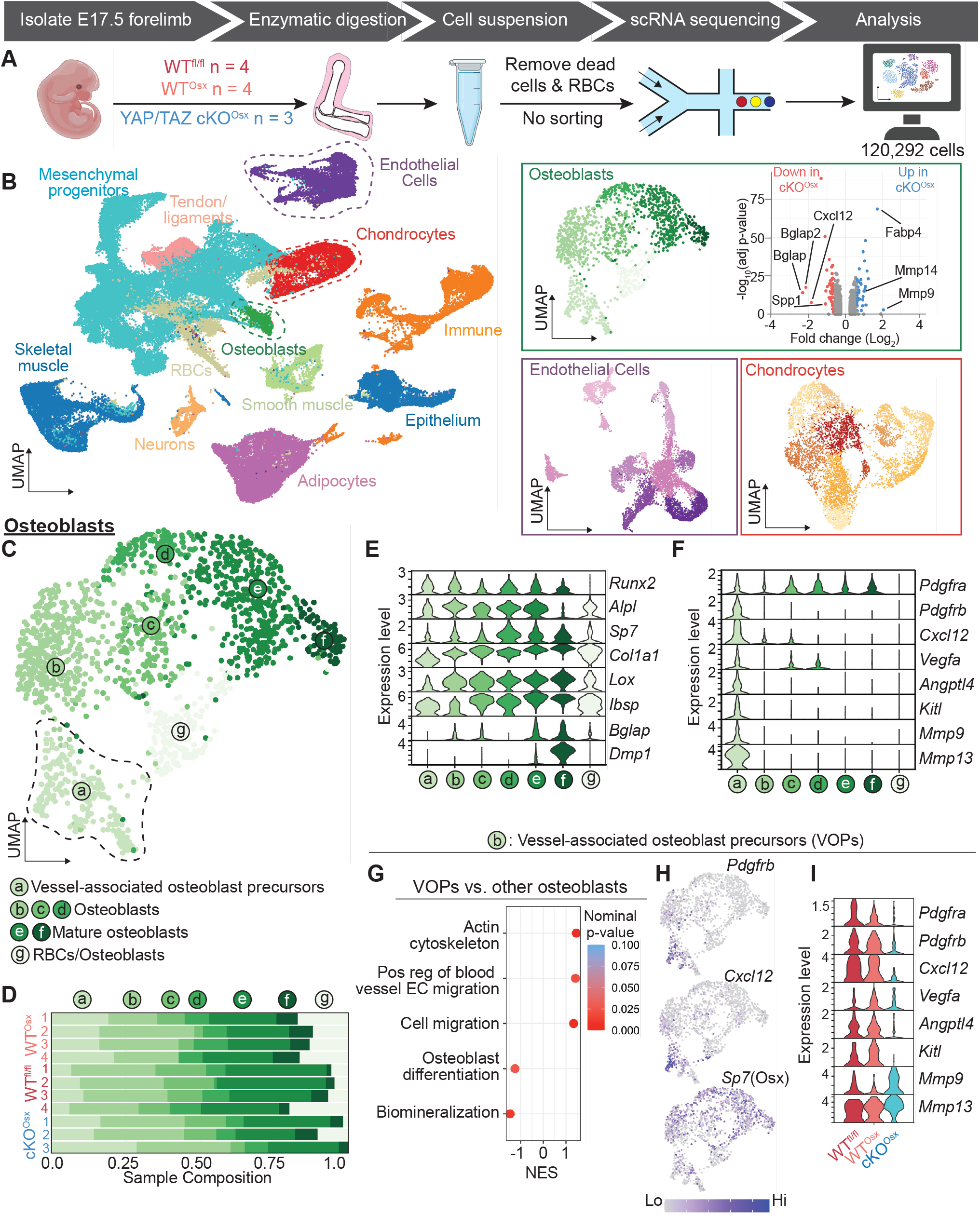
Identification of vessel-associated osteoblast precursors (VOPs) in fetal limbs by single-cell RNA sequencing. **(A)** Schematic of single-cell RNA sequencing experiment design. Fetal forelimbs, without autopod, from wildtype floxed (WT^fl^), wildtype Osx-Cre::GFP (WT^Osx^), and Osx-conditional YAP/TAZ knockout (YAP/TAZ cKO^Osx^) were digested at E17.5 for single-cell RNA sequencing (scRNA-seq; n = 3-4 per genotype). **(B)** Uniform manifold approximation and projection (UMAP) representation of the integrated scRNA-seq data, labelled by corresponding cell type and iterative UMAP representation of scRNA-seq data of selected cell type subsets. Volcano plot shows differentially expressed genes in the osteoblast cluster. **(C)** UMAP of iterative osteoblast-lineage clusters indicating six osteoblast lineage cell states (a-f) and one characterized by erythrocyte co-lysis (g). **(D)** Sample composition of osteoblast cell states across each independent embryo, on y-axis. **(E)** Osteoblast marker gene expression in the osteoblast cell states. **(F)** VOP marker gene expression in the osteoblast cell states. **(G)** Gene set enrichment analysis comparing VOPs against the other osteoblast cell states. **(H)** Gene plot showing *Pdgfrb, Cxcl12*, and *Sp7* (Osx) expression in osteoblast-lineage cells. **(I)** Gene expression in VOPs by genotype.

We identified a single cluster, of the 77 clusters, as osteoblast-lineage cells based on gene expression patterns, including *Runx2, Alpl, Osx (Sp7), Col1a1, Lox, Spp1, Ibsp, Bglap, and Dmp1* (**Fig. 3C-E, Fig. S7-8**). Osx-conditional YAP/TAZ deletion significantly altered gene expression in osteoblasts, reducing expression of mineralization genes (*Spp1, Bglap, Bglap2*) and increasing expression of matrix metalloproteinases (*Mmp9, Mmp14*) and adipocyte genes (*Fabp4*) (Fig. 3B). One of the most-downregulated genes in osteoblasts was the chemokine, *Cxcl12*, which mediates mechanoregulation of bone formation, angiogenesis, and hematopoiesis (*14*). Therefore, we sought to identify which of the osteoblastic cell states expressed these genes. We identified seven osteoblastic cell states by Louvain clustering, including one identified as osteoblasts that had been co-lysed with erythrocytes, based on expression of hemoglobin genes (cluster/state g; Fig. 3C-E). YAP/TAZ deletion did not significantly alter total osteoblast number (Fig. S9) or osteoblastic cell state distribution (**Fig. 3D**). However, only one osteoblast cell state (state a), was enriched for *Cxcl12*.

Osteoblast cell state ‘a’ featured the transcriptional profile of vessel-associated osteoblast precursors (VOPs; Fig. 3F). Consistent with prior lineage-tracing and conditional knockout studies defining the VOPs (*15, 16*), these cells were *Osx*^*+*^, *Runx2*^*+*^, *Pdgfra*^*+*^, *Pdgfrb*^*+*^ and *Kitl*^*+*^. The VOPs were also enriched for *Vegfa, Angptl4*, and *Mmp9/13*. Gene set-enrichment analysis showed that, compared to other osteoblast-lineage cells, the VOPs were significantly enriched for gene sets associated with cytoskeletal dynamics, positive regulation of blood vessel endothelial cell migration, and cell migration (Fig. 3G). Relative to VOPs, the other osteoblastic cell states were enriched for gene sets associated with biomineralization and osteogenesis (Fig. 3G). We next evaluated the effects of YAP/TAZ deletion on gene expression in VOPs. YAP/TAZ deletion significantly reduced VOP expression of *Cxcl12* (Fig. 3H,I, Fig. S8-9). YAP/TAZ deletion also reduced VOP expression of the angiogenic gene, *Angptl4* (but not *Vegfa*), *Kitl* (SCF), and increased expression of matrix remodeling genes, *Mmp9* and *Mmp13* (Fig. 3I, Fig. S8). Together, these data identify a particular effect of YAP/TAZ deletion on a specialized population of osteoblast precursors that exhibit signatures of vessel-stromal cells and express angiogenic chemokines and growth factors.

### YAP and TAZ mediate Vessel associated osteoblast precursor (VOP)-endothelial cell crosstalk, blood vessel integrity, and hypertrophic cartilage remodeling

During limb development, VOPs co-mobilize with blood vessels and regulate angiogenesis and vascular integrity as pericytes (*15, 16*). As in isolated single-cells, deletion of YAP and TAZ from Osx-expressing cells decreased the abundance of *Cxcl12* mRNA in the primary ossification center adjacent to invading blood vessels (Fig. 4A). Bone-resident endothelial cells represent a unique subset of endothelial cells with particular structure and function. There are two major subtypes of bone endothelium: type H and type L (*2*). Type H blood vessels express high CD31 (*Pecam1*) and Endomucin (*Emcn*) and support new bone formation (*2*). Type L blood vessels express low CD31 and Endomucin and form the sinusoidal vasculature in the bone marrow (*2*). We identified 10 endothelial cell states in the limb by iterative clustering of the endothelial cluster. Known markers identified two cell states as putative bone-resident type H endothelial cells (Fig. 4B,C, Fig. S11).

**Figure 4.**
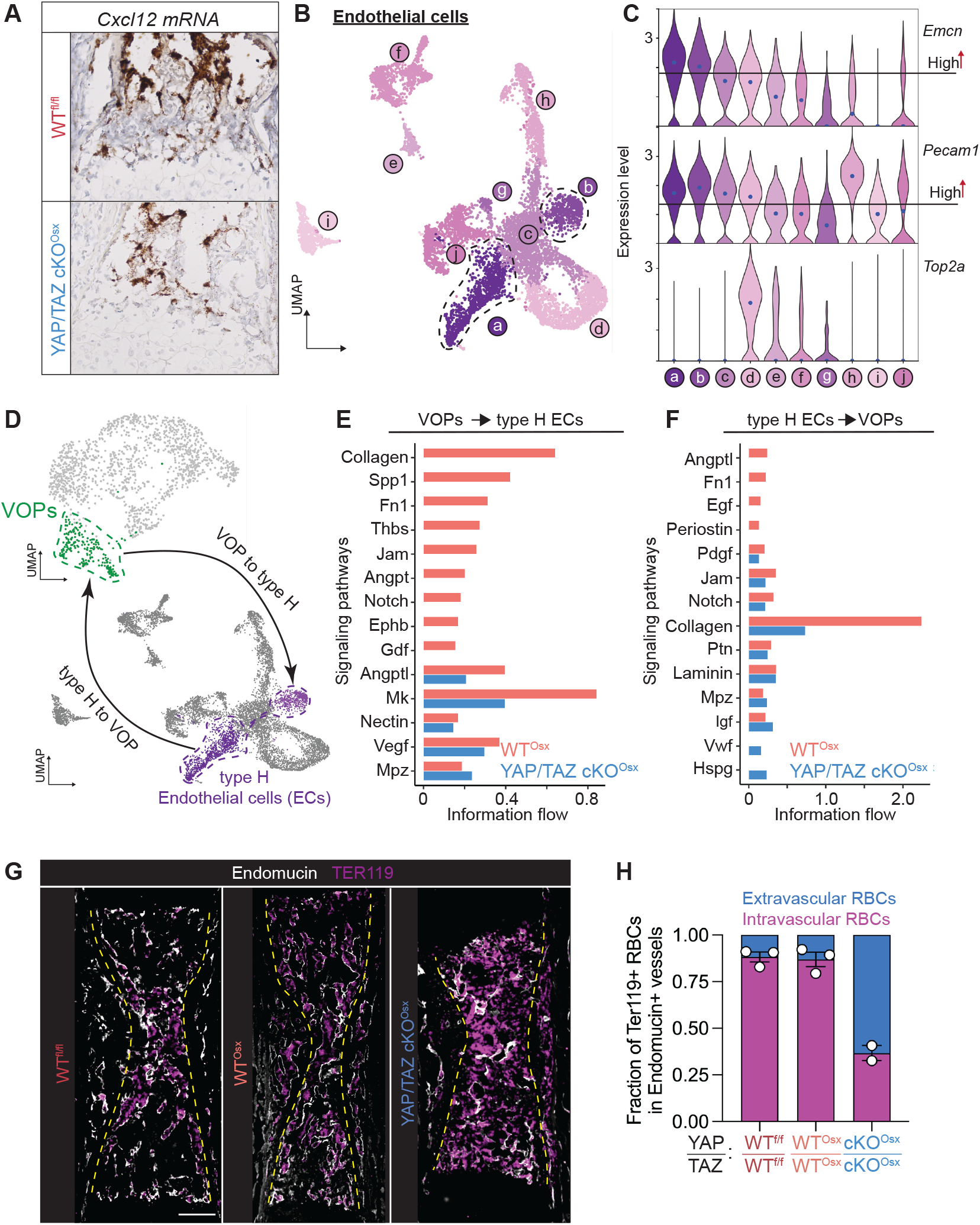
YAP/TAZ mediate vessel-associated osteoblast precursor(VOP)-endothelial cell crosstalk and vessel barrier function of primary ossification center blood vessels. **(A)**RNAscope *in situ* hybridization of *Cxcl12* mRNA in E17.5 growth plates (n = 1-2). (**B**) UMAP of iterative endothelial cell clusters, indicating ten endothelial cell states. (**C**) Gene expression analysis of Endomucin (*Emcn*) and CD31 (*Pecam1*), identifying endothelial states ‘a’ and ‘b’ as most-likely to represent EMCN-high/CD31-high (type H) bone-resident endothelial cells. **(D)** Representative crosstalk between cell states. **(E)** Occurrence of statistically-significant crosstalk pathways ranked by differences in information flow in the inferred networks between WT^osx^ and YAP/TAZ cKO^osx^. (**F**) Occurrence of statistically-significant crosstalk pathways ranked by differences in information flow in the inferred networks between WT^osx^ and YAP/TAZ cKO^osx^. **(G)** Immunofluorescent staining of erythrocyte-marker, Ter119, and blood vessel marker, Endomucin, in WT^fl/fl^, WT^osx^, and YAP/TAZ cKO^osx^ E17.5 forelimbs, to indicate extravascular red blood cells. **(H)** Quantification of the fraction of intravascular vs. extravascular erythrocytes. Scale bars: 200 μm. RBC: Red blood cells.

We next asked whether Osx-conditional YAP/TAZ deletion disrupted cell-cell communication between VOPs and endothelial cell states a and b (Fig. 4D, Fig. S11). CellChat analysis of cell-cell communication (*17*) revealed that YAP/TAZ deletion impaired both overall VOP-to-endothelial (Fig. 4E, Fig. S12-13) and endothelial-to-VOP crosstalk (Fig. 4F, Fig. S12-13). Functional crosstalk between vessel-stromal pericytes and endothelial cells maintains vascular barrier integrity (*18*). Supporting a role for YAP/TAZ in functional crosstalk, Osx-conditional YAP/TAZ deletion disrupted vascular barrier integrity, causing substantial hemorrhage in the primary ossification center, indicated by abundant extravascular TER119^+^ erythrocytes (Fig. 4G,H).

In embryonic bone, VOPs spatially associate with type H vessels. We next asked whether YAP and TAZ mediated the spatial coupling of Osx-expressing precursors with endothelial cells. YAP/TAZ deletion significantly increased the distance from a given Osx::GFP^+^ cell to its nearest Endomucin^+^ blood vessel (**Fig. 5A-B, Fig. S14**) and significantly altered the proximity distribution in the primary ossification center (Fig. 5C, D). YAP/TAZ deletion doubled the mean (± standard deviation) distance of Osx::GFP+ cells to their nearest blood vessel from 17.4±0.6 μm (WT^Osx^) to 36.1±8.8 μm (cKO^Osx^).

**Figure 5.**
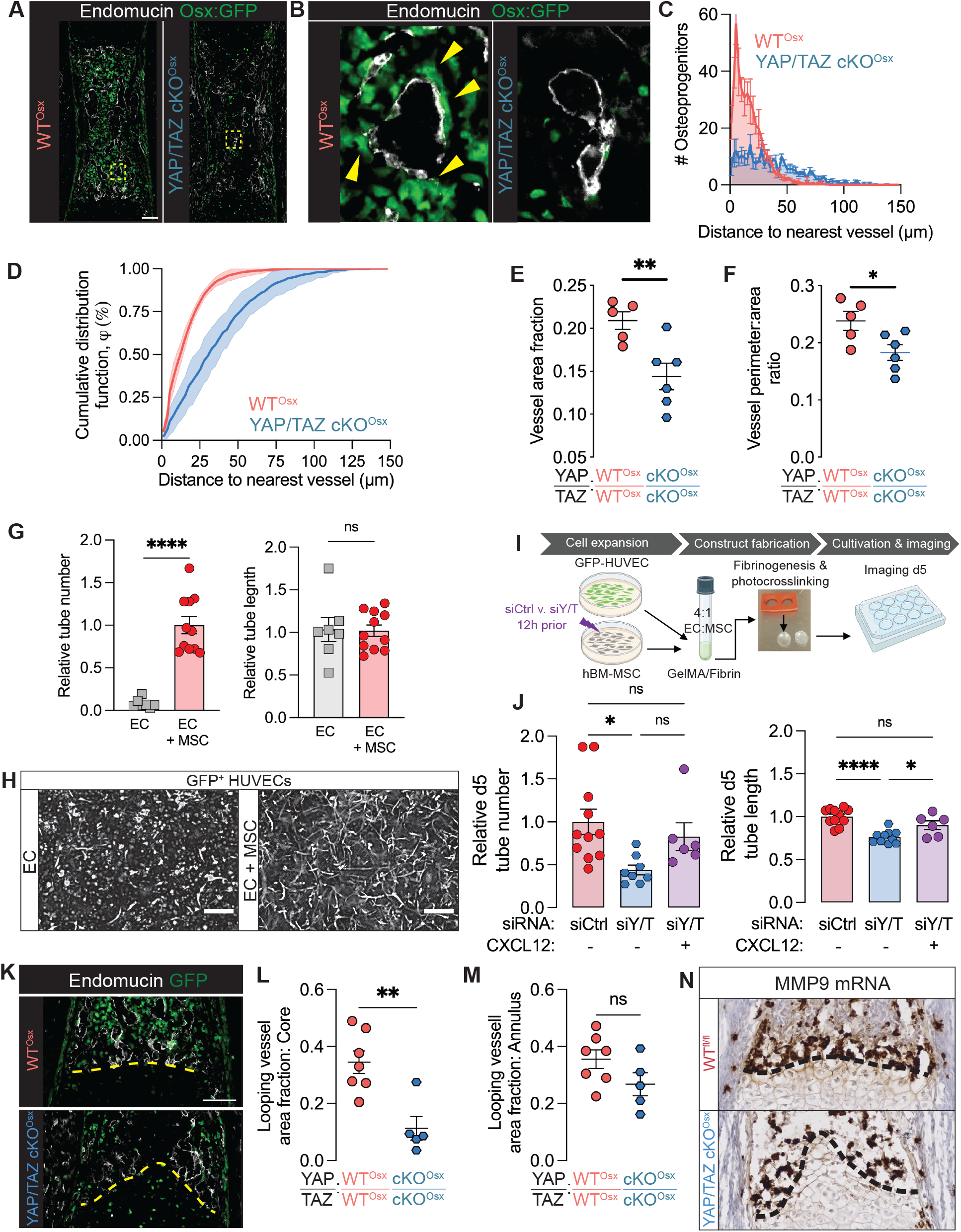
YAP/TAZ-Cxcl12 signaling spatially couples vessel-associated osteoblast precursors (VOPs) to angiogenesis in fetal bone. (**A**) Representative images of Endomucin immunostains in E17.5 humeri (n=5-6). **(B)** Selected magnified image of (A.). Yellow arrows indicate blood vessel-adjacent Osx::GFP^+^ cells. (**C**) Histogram of the distance from each Osx::GFP^+^ cell to the nearest Endomucin^+^ vessel. (**D**) Cumulative distribution function of the distance from each Osx::GFP^+^ cell to the nearest Endomucin^+^ vessel. Indicates the percentage of Osx::GFP^+^ cells within a given distance from a blood vessel. Error bars are SD. (**E**) Quantification of Endomucin-labelled blood vessel area relative to total area in the primary ossification center. (**F**)Measurement of average vessel size, quantified by cross-sectional perimeter to area ratio. (**G**) Quantification of relative tube number and tube length in GFP+ HUVEC 3D co-culture with and without hMSCs. (**H**) Representative images of HUVEC tube morphology after 5 days. Scale bar: 500 μm. (**I**) Schematic of 3D co-culture HUVEC + hMSC experiment with siRNA-mediated YAP/TAZ depletion from hMSCs and exogenous recombinant CXCL12 (100 ng/ml) rescue. (**J**) Quantification of relative tube number and tube length at day 5 following YAP/TAZ depletion in hMSCs and CXCL12 rescue.(**K**) Representative images of anastomotic looping vessels in E17.5 humeri. Yellow line indicates the chondro-osseous junction (**L**,**M**) Quantification of looping vessels in the core and annulus regions adjacent to the growth plate. (**N**)RNA scope in situ hybridization of *Mmp9* mRNA at the chondro-osseous junction (n=2). Scale bars: 100 μm, except 500 μm in panel H. Error bars: SEM unless stated otherwise. ‘*’: p<0.05; ‘**’: p<0.01; ‘***’: p<0.001; ‘****’: p<0.0001.

This shift in proximity distribution can be potentially explained by 1) a reduced attraction of Osx::GFP^+^ cells to endothelial cells, 2) a reduced number of Osx::GFP^+^ cells, or 3) impaired vessel distribution. To control for these factors, we simulated spatial Osx::GFP^+^ cell position and number within each blood vessel map. We determined that the proximity distribution was determined by the vascular morphology within the primary ossification center (Fig. S14). Consistently, Osx-conditional YAP/TAZ deletion significantly decreased vessel area fraction and vessel perimeter-to-area ratio (Fig. 5E,F), indicating a small number of larger vessels in the cKO^Osx^ mice. Together, these data suggest that Osx-conditional YAP/TAZ deletion impaired osteoblast precursor-endothelial cell spatial coupling by altering vascular morphogenesis in the embryonic bone.

Based on these functional defects in vascular morphogenesis and single cell RNA-seq identification of Cxcl12, we hypothesized that YAP/TAZ signaling promotes vascular morphogenesis via CXCL12. To test this directly in human cells, we used a 3D in vitro system, co-culturing GFP-expressing human umbilical vein endothelial cells (HUVECs) and human bone marrow stromal cells (hMSCs) to form tubular vascular networks (Fig. 5G,H). hMSCs were necessary for neovessel stability and promoted sustainable vascular network formation over 5 days (Fig. 5G,H). Tubular network formation was significantly impaired by YAP/TAZ depletion from hMSCs (Fig 5I,J), but was rescued by addition of recombinant CXCL12 (SDF-1) (Fig. 5J). These data support a functional role of CXCL12 in YAP/TAZ-mediated angiogenesis.

Endothelial cells also mediate cartilage matrix degradation in endochondral development, both directly and indirectly. Directly, anastomotic looping vessels at the chondro-osseous junction produce matrix metalloproteinases (MMPs, principally MMP9), which degrade cartilage matrix (*3*). Osx-conditional YAP/TAZ deletion disrupted the distribution of these vessels across the cartilage septum (Fig. 5K). Therefore, we quantified the density of anastomotic looping vessels in two regions of interest at the chondro-osseous junction: a core region and an outer annular region (Fig. 5L-M, Fig. S15). Osx-conditional YAP/TAZ deletion significantly altered anastomotic looping vessel density in the core, but not the outer annulus region. Further, *Mmp9*-expressing cells were reduced in this region (Fig. 5N). Together, these data suggest that osteoblast precursors spatially regulate vascular invasion and looping vessel morphogenesis at the chondro-osseous junction for growth plate remodeling. Indirectly, these growth plate-adjacent vessels also import and support cartilage-degrading cells, including septoclasts and osteoclasts (*19, 20*). Together, these data suggest that osteoblast precursors spatially regulate vascular invasion and looping vessel morphogenesis at the chondro-osseous junction for growth plate remodeling.

Osx-Cre also targets late hypertrophic chondrocytes (*21*). Therefore, to assess chondrocytic YAP/TAZ contributions to growth plate morphogenesis, we explanted limbs for *ex vivo* culture in the absence of a connected vascular supply. Hindlimbs were isolated at E15.5 and cultured for 6 days, as described previously (*22*) (Fig. 6A). In contrast to the intact developing limbs, Osx-conditional YAP/TAZ deletion did not impair transverse cartilage septum morphology (**Fig. 6B**), suggesting that continued vessel invasion is responsible for hypertrophic cartilage remodeling at the chondro-osseous junction during endochondral ossification.

**Figure 6.**
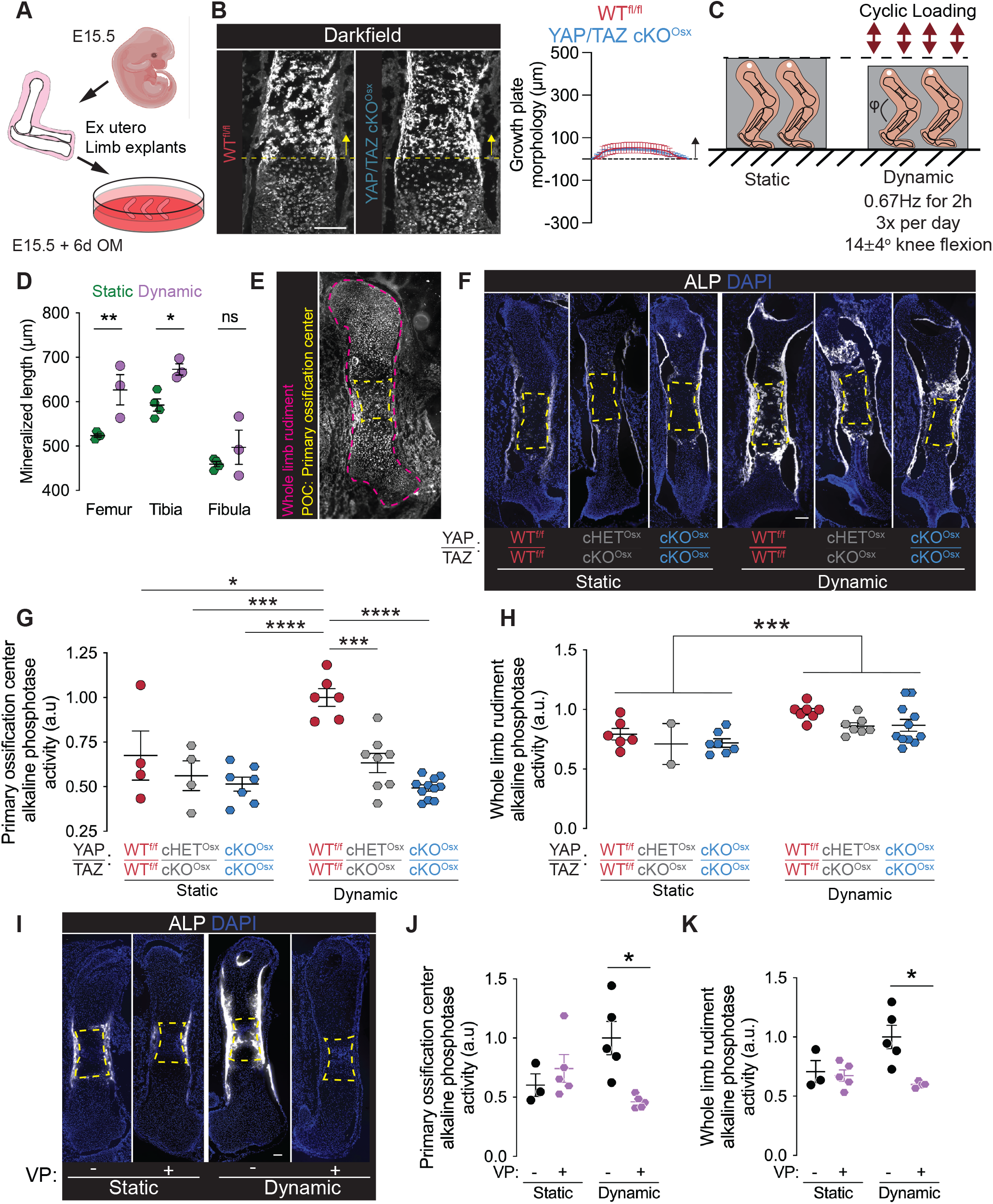
YAP/TAZ mediate mechanoregulation of fetal bone formation *ex vivo*. (**A**) Schematic of explant culture. **(B)** Growth plate morphogenesis in E15.5 explanted hindlimbs after 6 days static culture and quantification of growth plate morphology. (**C**) Schematic of bioreactor-based mechanical loading of explanted fetal hindlimbs. (**D**) Quantification of mineralized rudiment length in C57Bl6 in static and dynamic conditions after 5 days culture by optical projection tomography(**E**) Regions of interest indicating whole rudiment (magenta) and primary ossification center (POC, yellow). (**F**) Representative images of alkaline phosphatase (ALP) activity stains. (**G, H**) Quantification of ALP activity stain in regions of interest. (**L**) Representative images of ALP activity stains. (**J**,**K**) Quantification of ALP activity stain in regions of interest. Scale bars: 100 μm. Error bars: SEM.. ‘*’: p<0.05; ‘**’: p<0.01; ‘***’: p<0.001; ‘****’: p<0.0001

### YAP and TAZ mediate mechanoregulation of ex vivo bone formation

Endochondral ossification is regulated by mechanical cues, as seen in fetal akinesia (*5*), but the underlying mechanisms are unknown. Our findings here establish critical roles for YAP and TAZ in fetal bone development. Given their activation by mechanical cues, these data position YAP and TAZ as potential mediators of mechanoregulation of development. To test this, we performed bioreactor-based *ex vivo* mechanical stimulation of explanted E15.5 embryonic mouse hindlimbs (Fig. 6C). *Ex vivo* mechanical loading increased rudiment mineralization in the hindlimb (Fig. 6D). Osx-conditional YAP/TAZ deletion abrogated load-induced ALP activity in the primary ossification center (Fig. 6E-G), but not the whole rudiment, in which loading elevated ALP activity regardless of genotype (Fig. 6F,H). To account for contributions of YAP/TAZ deletion to developmental history, we used an orthogonal approach of acute pharmacologic YAP/TAZ inhibition in wild type embryos using Verteporfin (*9*). Unlike Osterix-conditional genetic deletion, global YAP/TAZ inhibition abrogated load-induced ALP activity, both in the primary ossification center (Fig. 6I,J) and in the whole limb rudiment (Fig. 6I,K). Thus, Osx-conditional YAP/TAZ deletion, which predominantly altered endochondral bone development in the primary ossification center, also prevented load-induced ALP activity in this region, while acute global inhibition prevented load-induced activity throughout the skeletal rudiment. Together, these data implicate YAP/TAZ mechanosignaling in fetal movement-directed bone development.

## Discussion

Here, we establish functional roles of the vessel-associated osteoblast precursors (VOPs) that co-mobilize with blood vessels in fetal endochondral bone development. The transcriptional regulators YAP and TAZ mediate VOP mobilization and function for primary ossification center initiation and expansion. Mechanistically, YAP/TAZ activity in Osterix-expressing cells spatially couples VOPs to blood vessels, maintains vascular barrier integrity, coordinates vascular morphogenesis, via CXCL12, to regulate growth plate remodeling, and mediates mechanoregulation of embryonic bone formation. Together, our findings establish a model of bone development in which VOPs direct angiogenesis, and subsequent endochondral bone development, through YAP/TAZ-*Cxcl12* signaling.

Osteoblast precursors invade the cartilage template prior to endochondral ossification. These cells express the osteogenic transcription factor, Osterix, and mobilize before their differentiation into mature osteoblasts (*1*). Here, we show that Osterix-conditional YAP/TAZ deletion impaired osteoblast precursor mobilization and subsequent endochondral bone formation. Limb development features both endochondral ossification in the primary ossification center and intramembranous bone formation in the bone collar (*23*). Both are accomplished by osteoblasts, but while the former requires precursor mobilization, the latter occurs by local osteoblast differentiation from perichondrium-resident progenitors (*24*).We found that Osterix-conditional YAP/TAZ deletion significantly reduced both total cell density and Osx::GFP^+^ cell density in the primary ossification center, but only moderately reduced either in the bone collar. Previously, we found that YAP and TAZ are required for persistent cell migration via a feedback loop that maintains dynamic cytoskeletal equilibrium (*8, 25*). Therefore, we performed actin cytoskeleton staining in collagen reporter mice that mark the three primary endochondral cells (chondrocytes, hypertrophic chondrocytes, and osteoblasts), and identified the emergence of osteoblast precursors in the primary ossification center. This osteoblast precursor mobilization was both delayed and reduced by YAP/TAZ deletion. Consistently, YAP/TAZ deletion significantly reduced osteoblast activity, measured by alkaline phosphatase activity, in the primary ossification center, but not in the bone collar. Together, these data show that YAP and TAZ mediate osteoblast precursor mobilization into the primary ossification center and mediate endochondral ossification and implicate distinct functions in endochondral vs. intramembranous ossification.

Endochondral osteoblast precursors co-mobilize with invading blood vessels as vessel-associated osteoblast precursors (VOPs). Here, we show that Osterix-conditional YAP/TAZ deletion impaired osteoblast-lineage cell gene expression, and particularly disrupted angiogenic gene expression in VOPs. Single-cell RNA sequencing confirmed YAP/TAZ-regulated genes in osteoblast-lineage cells that we previously identified in bone and tendon cells (i.e., *Bglap, Bglap2, Mmp9, Mmp14*) (*9, 26*–*28*). Notably, YAP/TAZ deletion also significantly reduced *Cxcl12* mRNA in osteoblast-lineage cells. Among fetal osteoblast-lineage cells, *Cxcl12* was primarily expressed by a single sub-cluster. This *Runx2*^*+*^, *Pdgfra*^*+*^, *Osx*^*+*^ population was uniquely marked by expression of *Pdgfrb*^*+*^ *and Kitl*^*+*^, and additionally moderate *Lepr*^*+*^ (*29*) (Fig. S8), which recent lineage tracing studies identify as vessel-associated osteoblast precursors (VOP) (*15, 16*). YAP/TAZ deletion particularly altered gene expression in VOP cells, relative to other osteoblast-lineage cells. In the VOP cluster, YAP/TAZ deletion reduced expression of *Cxcl12* as well as *Angptl4*, and increased expression of *Mmp9* and *Mmp13. Cxcl12* regulation in VOPs is consistent with recent reports showing that CXCL12 mediates mechanoregulation of vascularized bone formation (*30*–*32*). *Cxcl12* is an angiogenic and myeloid cell-chemotactic chemokine (*14*) that is expressed by a variety of cell types in adult bone marrow, especially CXCL12-abudant reticular (CAR) cells (*33*). However, multiple recent studies show no evidence of CAR cells in embryonic or fetal bone marrow (*34, 35*). Confirming a functional role for CXCL12 signaling in YAP/TAZ-induced mesenchymal-endothelial crosstalk, we found that recombinant CXCL12 treatment rescued effects of human mesenchymal cell YAP/TAZ depletion on 3D human endothelial cell network formation *in vitro*. Together, this establishes an angiogenic YAP/TAZ-CXCL12 axis in vessel-associated osteoblast precursors that couples osteoblast precursor mobilization to vessel morphogenesis, but whether YAP and TAZ regulate Cxcl12 by direct transcriptional activation or other means will require further research beyond the scope of the present study.

Vessel-associated osteoblast precursors regulate vascular function. Here, we show that Osterix-conditional YAP/TAZ deletion impaired vascular barrier function and vessel-mediated cartilage matrix remodeling. PDGFRβ^+^ vessel stromal cells functionally support blood vessel morphogenesis and vascular barrier function (*18*). YAP/TAZ deletion from Osx^+^/PDGFRβ^+^ vessel-associated osteoblast precursors (VOPs) reduced their spatial association with blood vessels, which spatial variation analysis showed to be driven by dysregulated vessel morphogenesis. These defects in vascular morphogenesis caused two primary functional defects. First, Osx-conditional YAP/TAZ deletion impaired vascular barrier function, possibly through reduced CXCL12 (*36*), leading to vessel leakiness in the primary ossification center, indicated by extensive erythrocyte extravasation. Second, Osx-conditional YAP/TAZ deletion impaired hypertrophic cartilage remodeling. Hypertrophic matrix removal is mediated both by cell-autonomous chondrocytic matrix lysis and by degradation at the chondro-osseous junction by cartilage-degrading capillaries and other blood vessel-recruited cells (*3, 19*). Osx-conditional YAP/TAZ deletion severely impaired cartilage remodeling, lengthening the hypertrophic zone and disrupting the characteristic uniformity of the transverse cartilage septum. Osx-Cre is expressed in both hypertrophic chondrocytes and osteoblast-lineage cells, suggesting both chondrocyte-autonomous and non-autonomous remodeling mechanisms. In a comparable study, Col2-Cre-conditional YAP/TAZ deletion elongated the hypertrophic zone but maintained a flat transverse cartilage septum (*37*), suggesting that the hypertrophic elongation is hypertrophic chondrocyte-autonomous. In contrast, the conical shape of the cartilage septum correlates with the organization of early vessel invasion (*38*) and with the disruption of *Mmp9*-expressing cell organization along the chondro-osseous junction. Further, our *ex vivo* limb culture experiments, in which limbs resected from their vascular supply after initiation of the primary ossification center, failed to produce a conical chondro-osseous junction regardless of YAP/TAZ deletion status, suggesting a role for vessel-associated cells in remodeling of the hypertrophic cartilage. Future study will be required to dissect the role of YAP/TAZ signaling crosstalk in the cartilage-degrading capillaries (*19, 20, 39*).

Lastly, fetal muscle contractions induce mechanical stimuli that are required for proper bone development. Clinically, fetal akinesia (i.e., insufficient movement from caused by low amniotic fluid volume or other), or impaired muscle development, can cause skeletal disorders such as hip dysplasia, arthrogryposis, and impaired bone development (*5*). We developed a bioreactor system for *ex vivo* mechanical stimulation of explanted fetal limbs (*22*). Pharmacologic inhibition shows that the ion channel Trpv4(*40*) mediates skeletal mechanosensing, but the mechanotransductive signaling mediators are unknown. YAP and TAZ mediate mechanotransduction in other embryonic tissues (*41*), and are dysregulated in a mouse model of fetal akinesia(*42*). Here, we show that YAP and TAZ mediate mechanical load-induced fetal bone formation ex vivo using local orthogonal genetic and global pharmacologic approaches.

Together, these data identify YAP and TAZ as mechanoresponsive transcriptional regulators that couple osteoblast precursor mobilization to vessel morphogenesis and mechano-regulated bone development.

## Limitations

This study has several limitations. First, our study focused only on wild type and double homozygous knockout mice, and does not inform the relative roles of YAP vs. TAZ. We chose this based on our prior study identifying YAP and TAZ as combinatorial regulators of postnatal bone development, with perinatal lethality only in double homozygous knockout mice (*9*). Future study will be required to dissect the respective roles of YAP or TAZ in embryonic bone. Second, we selected Osterix-Cre based on the predominant distribution of YAP/TAZ expression in skeletal-lineage cells (*9*). The Osterix-Cre driver has been observed to exhibit a skeletal phenotype in the absence of floxed alleles (*43, 44*). However, the magnitude of the effect on skeletal development is determined by the genetic background. We controlled for this using both floxed-alone and Cre-containing controls. As we described previously(*26, 27*), the Osx-Cre did not interfere with the effects of YAP/TAZ signaling in these mice. Lastly, our work identifies VOP-mediated vessel morphogenesis as a critical regulator of embryonic bone development, but our bioreactor loading experiments that identify YAP and TAZ as mechanotransducers in embryonic bone required limb explants that are disconnected from the native vascular supply. Future study to enable modulation of *in utero* mechanical stimulation will be required to directly test the roles of YAP and TAZ in mechanoregulation of skeletal morphogenesis *in vivo*.

## Methods and Materials

### Animals

We conditionally deleted YAP and TAZ from Osx1-expressing cells (YAP^fl/fl^;TAZ^fl/fl^;Osx1-GFP::Cre) comparison to littermate wild type (YAP^fl/fl^;TAZ^fl/fl^) and Osx1-GFP::Cre wild type (YAP^WT/WT^; TAZ^WT/WT^;Osx1-GFP::Cre) controls with a mixed C57BL/6 background (*9, 21*). Additionally, we used a triple transgenic fluorescent reporter model which co-expresses fluorescent proteins under the control of promoters for three collagen genes: Collagen II, marked by CFP (Col2-CFP); Collagen X, by mCherry (ColX-mCherry); and the 3.6kb fragment of the Collagen1a1 promoter, by YPF (Col1-YFP). The Col1-YFP is expressed in both immature and committed osteoblasts. To generate embryos by timed pregnancies, breeding pairs were housed together for a single night and pregnancy was determined by visual inspection and increased female weight gain over the course of the pregnancy. To generate YAP/TAZ cKO^osx^/WT^fl/fl^ littermate embryos, a YAP/TAZ cKO^osx^ stud male was housed with a WT^fl/fl^ female overnight, which was never exposed to doxycycline. To generate WT^osx^ embryos, a WT^osx^ stud male was housed with a YAP^WT/WT^;TAZ^WT/WT^ female overnight. Pregnant mice were euthanized by CO^2^ and secondary cervical dislocation and embryos were harvested at 15.5 days (E15.5) and 17.5 days (E17.5) post-conception. Mice were genotyped by an external service (Transnetyx, Cordova, TN, USA). All procedures were performed with IACUC approval (protocol: 806281).

### Histology and immunofluorescence

For cryohistological analysis, embryos were fixed for 24-48 hours in 4% PFA (Fisher) in water at 4°C. Only samples fixed for similar times were compared. Following fixation, samples were saturated in 30% sucrose/PBS at 4°C and embedded in OCT (Fisher). Samples embedded in OCT were stored at -20°C until further use. 7 μm sections were generated by cryohistology (NX70 cryostat Thermo Fisher Scientific), using methods previously described (*45*). For immunocytochemistry, tissue sections were washed 3x 5 mins with PBS. Then, samples were permeabilized and blocked for 1hr at RT with blocking buffer (PBS/ 5% goat serum/ 0.3% Triton X-100 (MP Biomedical). Following this, cells were stained with primary antibodies overnight at 4°C in a dilution buffer (PBS/ 0.01%BSA/ 0.3% Triton): YAP (1:100, D8H1X #14074 Cell Signaling Technology), TAZ (1:100, 72804S Cell Signaling Technology), Collagen-10 (1:100, ab58632 Abcam), GFP (1:1000, ab13970 Abcam), Endomucin (1:50, sc-65495 Santa Cruz), Ter119-APC (1:100, BioLegend 116211). The next day, samples were washed 3x 5 mins with PBS and then stained with corresponding secondary antibodies (1:1000; 4414S Cell Signaling Technology, 8889S Cell Signaling Technology, A21247 Life Tech). F-actin was stained with Alexa Fluor-conjugated Phalloidin (1:40, Invitrogen Thermo Fisher Scientific) for 2hr. Alkaline Phosphatase was stained with the Vector blue Alkaline phosphatase substrate kit (Vector SK-5300), per manufacturer’s instructions. Sections were stained with Calcein blue, according to Dyment *JOVE* 2016(*45*). Following staining, samples were mounted in Prolong Diamond (Invitrogen Thermo Fisher Scientific) or 50% glycerol/PBS. RNAscope was performed according to Advanced Cell Diagnostics manufacturer’s instructions.

### Histological imaging and analysis

Fluorescent images were captured with an Axioscan (Carl Zeiss Microscopy Deutschland GmbH). Data acquisition was performed with ZEN imaging suite (Zeiss). Image analysis (cell density, morphometric measurements, fluorescent intensity, and coordinate values) were performed with ImageJ (FIJI). Primary ossification center, bone collar, and hypertrophic zone regions of interest were determined manually by by darkfield tissue morphology and additionally informed by relevant immunofluorescent stains (e.g. Collagen-10). ROIs were managed with the ImageJ’s built-in ROI-Manager. The cell number within each ROI was determined manually with the Cell Counter Plugin in ImageJ and DAPI signal. The line plots of fluorescent intensity (Fig. 2B,C) were generated from the average intensity of 20-30 μm width bins orthogonal to the central axis along the bone rudiment. Each average fluorescent intensity measurement was normalized to the area in which that measurement was taken. For the Col2-CFP, Col10-RFP, Col1(3.6kb)-YFP line plot analysis, the area-normalized fluorescent intensity of each channel in a region with external to the tissue was subtracted from the area-normalized tissue intensity. To quantify the distance between Osx::GFP^+^ cells and Endomucin^+^ vessels (Fig. 5C,D), we generated maps of coordinates for GFP^+^ cell positions and vessel positions in each primary ossification center. We used R to determine the nearest vessel to each GFP^+^ cell position. Further, to evaluate different explanations for the proximity distributions, we computationally varied the spatial position of each Osx::GFP+ cell within the primary ossficiation center. To account for the contribution of Osx::GFP+ cell-endothelial cell attraction, we took the Endomucin+ vessel map from both WT and cKO samples and then randomly distributed the Osx:GFP+ cells within that sample (Fig S15). Thus, if Osx::GFP^+^ cell-endothelial cell attraction explained the proximity distribution, then randomizing the distribution of Osx::GFP+ cells should produce equivalent proximity distributions in both WT and cKO samples. To account for the contribution of Osx::GFP+ cell number, we took the Endomucin+ vessel map from both WT and cKO samples and then randomly distributed an equivalent number of Osx::GFP+ cells (Fig S15). Thus, if the reduced number of Osx::GFP^+^ cells explained the proximity distribution, then randomly distributing an equivalent number of Osx::GFP+ cells to both WT and cKO vessel maps should produce equivalent proximity distributions. To account for the contribution of vessel distribution, for any possible Osx::GFP^+^ cell position, we took the Endomucin^+^ vessel map from both WT and cKO samples and mapped the proximity distribution of all extravascular pixels in the primary ossficiation center (Fig S15). Looping vessels were calculated as Endomucin+ vessels within 50 μm of the chondro-osseous interface. For bioreactor analysis, during image quantification of regions of interest, samples were excluded if the whole limb rudiment was not intact or if the primary ossification center-bone collar interface was unclear.

### Tissue preparation for single-cell RNA sequencing

Timed pregnancies and mouse embryos were generated, as described above. For tissue isolation, embryonic forelimbs were isolated on ice and trimmed to removed excess and loose skin/soft tissue and the paw. Individual embryonic tails or yokesacs were placed aside for genotyping. A single-cell suspension was isolated as previously described (*13*), in the presence of the transcriptional inhibitor Actinomycin D (MP Biomedicals). Briefly, cells from individual samples were disassociated from the tissue with a digestion media including DMEM, fetal bovine serum (Sigma), Collagenase II (Gibco Thermo Fisher Scientific), Pronase (Sigma Aldrich), and Actinomycin D (2 μg/ml) for 75 minutes at 37°C. Following, cells were washed with PBS/0.01% BSA and passed through 70 μm cell strainers. Then, red blood cells were lysed with a Red Blood Cell Lysis kit (Miltenyi Biotec, 130-094-183), per manufacturer instructions. Dead cells were then removed with a Dead Cell Removal kit (Miltenyi Biotec 130-090-101) and MACS MS columns (Miltenyi Biotec), and a MiniMACS Magnetic Cell Separator (Miltenyi Biotec). Following dead cell removal, each sample was confirmed to have >95% viability. To allow time for genotyping, cells were then processed according to 10x Genomics ‘Methanol Fixation of Cells for Single-cell RNA Sequencing protocol, Rev D’. Cell suspensions were resuspended per 10x Genomics ‘Methanol Fixation of Cells for Single-cell RNA Sequencing protocol, Rev D’ and processed for sequencing.

### Single-cell RNA sequencing and analysis

Next-generation sequencing libraries were prepared using the 10x Genomics Chromium Single-cell 3’ Reagent kit v3 per manufacturer’s instructions. Libraries were uniquely indexed using the Chromium dual Index Kit, pooled, and sequenced on an Illumina NovaSeq 6000 sequencer in a paired-end, dual indexing run. Sequencing for each library targeted 77,000 mean reads per cell. Data was then processed using the Cell Ranger pipeline (10x Genomics, v.6.0.0) for demultiplexing and alignment of sequencing reads to the mm10 transcriptome and creation of feature-barcode matrices. For further analysis, we used Seurat v4.0. Cells with less than 600 genes, greater than 6000 genes, or those with greater than 5% mitochondrial reads were removed. Data from individual samples were LogNormalized. We identified 2000 variable features across each sample. Samples were integrated using the Seurat alignment method for data integration (*46*). Samples were scaled using ScaleData. Linear dimensional reduction was conducted using principal component analysis (PCA). The entire 120,292-cell dataset was projected onto two-dimensional space using UMAP, and the data were clustered using Louvain clustering. The analyses on the whole limb used 50 PCA dimensions and a resolution of 2.5. The FindAllMarkers function and canonical marker genes was used to identify major cell types. The 77 clusters identified by Louvain clustering were merged based on similar expression patterns into 12 distinct major cell populations. Osteoblasts constituted only 1 of the initial 77 clusters. These cells were subsetted and rerun through the above pipeline. The analysis of the osteoblasts used 15 PCA dimensions and a resolution of 0.5. The iterative clustering of the osteoblasts revealed 7 cell states, which were identified using the FindAllMarkers function and cell state specific markers determined in prior lineage tracing studies. For gene set enrichment analysis (GSEA), a pre-ranked list of the genes was constructed by fold change between comparisons and evaluated with GSEA v4.2.3. Significant inter-cluster gene expression differences between genotypes was determined by MAST with Bonferroni corrections. Each wildtype was separately compared by MAST with Bonferroni corrections to the Osx-conditional YAP/TAZ knockout samples. Additionally, endothelial cells and chondrocytes were subsetted and rerun through the above pipeline. The analysis of the endothelial cells used 15 PCA dimensions and a resolution of 0.2. Clusters which were mostly likely to contain type H endothelial cells were determined by their comparative expression of *Emcn* and *Pecam1*. The analysis of the chondrocytes used 8 PCA dimensions and a resolution of 0.5. Cell-cell communication analysis was done with the package CellChat (*17*), which accounts for ligand and receptor expression, as well as co-expression of genes that are agonistic and antagonistic to a particular communication pathway. Cell-cell communication analysis via CellChat was performed on the entire osteoblast and endothelial cell populations and specifically between vessel-associated osteoblast precursors and type H endothelial cells.

### 3D in vitro vascular network assay

GFP+ HUVECs (Angio-Proteomie) were expanded in human endothelial growth medium with 5% FBS (EGM-SF1; Angio-Proteomie, Boston, AM, USA) in cell culture flasks coated with 0.2% gelatin (Sigma Aldrich). hMSCs were cultured in RoosterNourish™-MSC-XF (both RoosterBio). For RNAi YAP and TAZ knock down – hMSCs were starved for 6h in DMEM without antibiotics before adding siRNA for YAP or TAZ and scramble control (Stealth RNAi Invitrogen Thermo Fisher Scientific) in OptiMEM with addition of Lipofectamine 3000 (both Thermo Fisher Scientific). GFP+ HUVECs and hMSCs were embedded (4:1; total final cell concentration 2 Mio cells/mL) in photocrosslinked gelatin-fibrin hydrogels (CELLINK) composed of 5% GelMA, 5 mg/ml fibrinogen and 0.2% LAP for photoinitiation and cultivated in vitro for up to 5 days with endothelial basal medium containing 0.1% FBS and 1% P/S (Thermo Fisher Scientific). For rescue experiment, 100 ng/mL recombinant CXCL12/SDF-1 (350-NS/CF; R&D Systems) was added to the medium. Images were taken at 5 days using a Keyence BZ-X800 Fluorescence Microscope (Keyence). All images were blinded for group and treatment. The tube number and mean tube length were analyzed using Fiji ImageJ. The area to be analyzed was defined as the full area of the hydrogel which was visible in the images. To correct for differences in analyzed area (ROI) dimension between the hydrogels, the tube number was normalized to the analyzed area (ROI). Finally, the normalized tube number and the absolute mean tube length were calculated relative to the respective scramble control. To establish the assay we performed two independent experiments: In experiment 1, we used two different MSC lines combined with two different HUVEC lines with 2-3 replicates per MSC/HUVEC combination and condition. Experiment 2 was used to replicate and strengthen the data with one MSC/HUVEC combination and 4 replicates for each condition (biological replicates n= 5; technical replicates total n= 7-11). For the knockout and rescue experiment, we combined the data from two independent experiments – experiment 1 established the knockout with one MSC/HUVEC combination and 4 replicates for each condition. In the second experiment, we used two different MSC lines combined with one HUVEC line testing 2-3 replicates per MSC/HUVEC combination and condition and performed the rescue experiment (biological replicates n= 2-3; technical replicates total n= 6-11).

### *Ex vivo* bioreactor experiments

We explanted hindlimbs at E15.5 and cultured them in either dynamically loaded or static conditions in osteogenic media containing: αMEM, 1% Pen/Strep, GlutaMAX, 100μM Ascorbic acid (Sigma Aldrich) for 5-6 days. Soft tissue was first removed from the hindlimbs. The limb explants were pinned at the hip to a polyurethane deformable foam (Sydney Heath & Son) and positioned in a mechanostimulation bioreactor (Ebers TC-3). Dynamic loading was applied at the top of the foam at 0.67 Hz for 2h, 3x/day, producing cyclic knee flexion to 14±4°, which mimics motion produced through prenatal muscle contractions. Static conditions used the same set up but without dynamic loading. We evaluated the effect of loading on hindlimb explants cultured under static or dynamic conditions for 5 days. After culture, the limbs were evaluated by Optical Projection Tomography (OPT). For OPT, hindlimbs were dehydrated and stained for cartilage and mineralized tissue using alcian blue and alizarin red S, as previously described(*47*). Stained and fixed limbs were embedded in agarose, dehydrated and cleared in a solution of BABB in preparation for 3D imaging using OPT, according to (*48*).Limbs were scanned under visible light to obtain 3D images of the alcian blue staining (cartilage) and under the Texas-red filter to obtain auto-fluorescent 3D images of the alizarin red S-stained region (the mineralized region). Scans were reconstructed using NRecon (SkyScan, Bruker microCT, 2011). For the genetic knockout, E15.5 embryos were generated for 3 rounds of bioreactor experiments, as described above. In prior studies, Osx-conditional YAP/TAZ deletion impaired post-natal bone development in an allele-dependent manner. To consider a moderate loss in mechanotransduction, here we evaluated WT^f/f^;WT^f/f^, YAP cHET^Osx^/TAZ cKO^Osx^, and YAP/TAZ cKO^Osx^ embryos. Following culture, samples were processed for cryohistology in the method described above. The fluorescent intensity of samples was normalized to average dynamic wildtype of their respective round to account for variations between rounds. Growth plate morphology was quantified in limb explants under the static condition. For pharmacologic inhibition, C57BL/6J E15.5 hindlimbs were isolated. Limbs were preprocessed for culture in the above way. Limbs were cultured under dynamic or static conditions in osteogenic media and in the presence of 5mM verteporfin or a vehicle control (DMSO).

### Statistical analysis

GraphPad Prism V.8 and V.9 were used for statistical analysis. We performed t-tests or 1-way ANVOA with post hoc Tukey’s HSD, where appropriate. In cases with multiple variables, we performed 2-way ANOVA with post hoc Sidek’s multiple comparisons. Single-cell RNA sequencing statistics are discussed in the associated methods section above. A p-value <0.05 was considered statistically significant. Sample sizes are indicated in the figure legends. Where possible data is displayed with the individual samples shown.

Asterisks represent significant differences with a p-value <0.05. Multiple asterisks represent lower p-values, as indicated in the figure legend. Sample sizes indicate measurements of individual embryos.

### Code availability

Analysis of the data presented here used published R packages available on GitHub. For analysis using published R packages, we used: R v4.1.0, Seurat v4.1.1, and CellChat v1.6.0.

### Schematics

The cartoons in Fig. 3A, 6A, and 6C were created using BioRender.com

## General

The authors would like to thank all members of the Boerckel lab for constructive discussions, Dr. Robert Tower (UTSW) for helpful suggestions, Dr. Farshid Guilak for providing protocols for the isolation of cells from embryonic forelimbs, and the Children’s Hospital of Philadelphia Center for Applied Genomics for their assistance with scRNA-seq sample processing.

## Author contributions

Conceptualization: JMC, AL, CP, NN, and JDB; Analysis: JMC, AL, CP, YM, MN, JDB; Funding Acquisition, Project Administration, and Supervision: JDB; Investigation: JMC, AL, CP, YM, JHK, MN; Methodology: AL, GS, LQ, RG, ND, NCN, JDB; Resources: NCN, ND, LQ, JDB; Writing: JMC, JDB; Review and Editing: all authors.

## Data and materials availability

All data associated with this study are present in the paper or the Supplementary Materials. Sequencing data will be deposited to GEO upon acceptance.

## Supplementary figures

**Figure S1.**
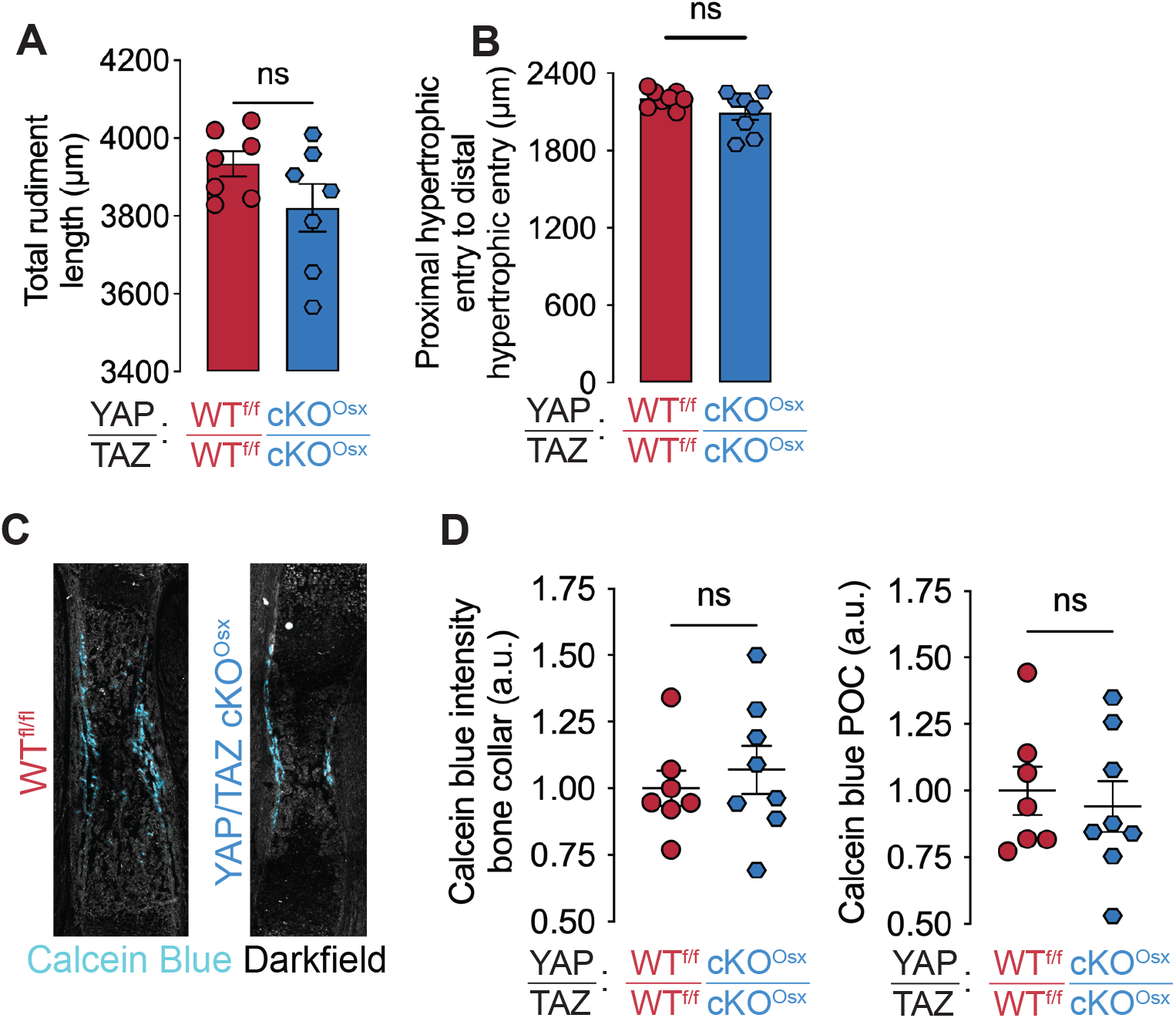
Additional data on the Osx-conditional YAP/TAZ knockout humeri at E17.5. (**A**) Length of the humerus rudiment at E17.5. (**B**) Distance between proliferating chondrocyte zones. i.e. the sum of both proximal and distal hypertrophic zones and primary ossification center at E17.5. (**C**) Calcein blue staining of E17.5 humerus. (**D**) Quantification of Calcein blue images in the bone collar and primary ossification center (POC).

**Figure S2.**
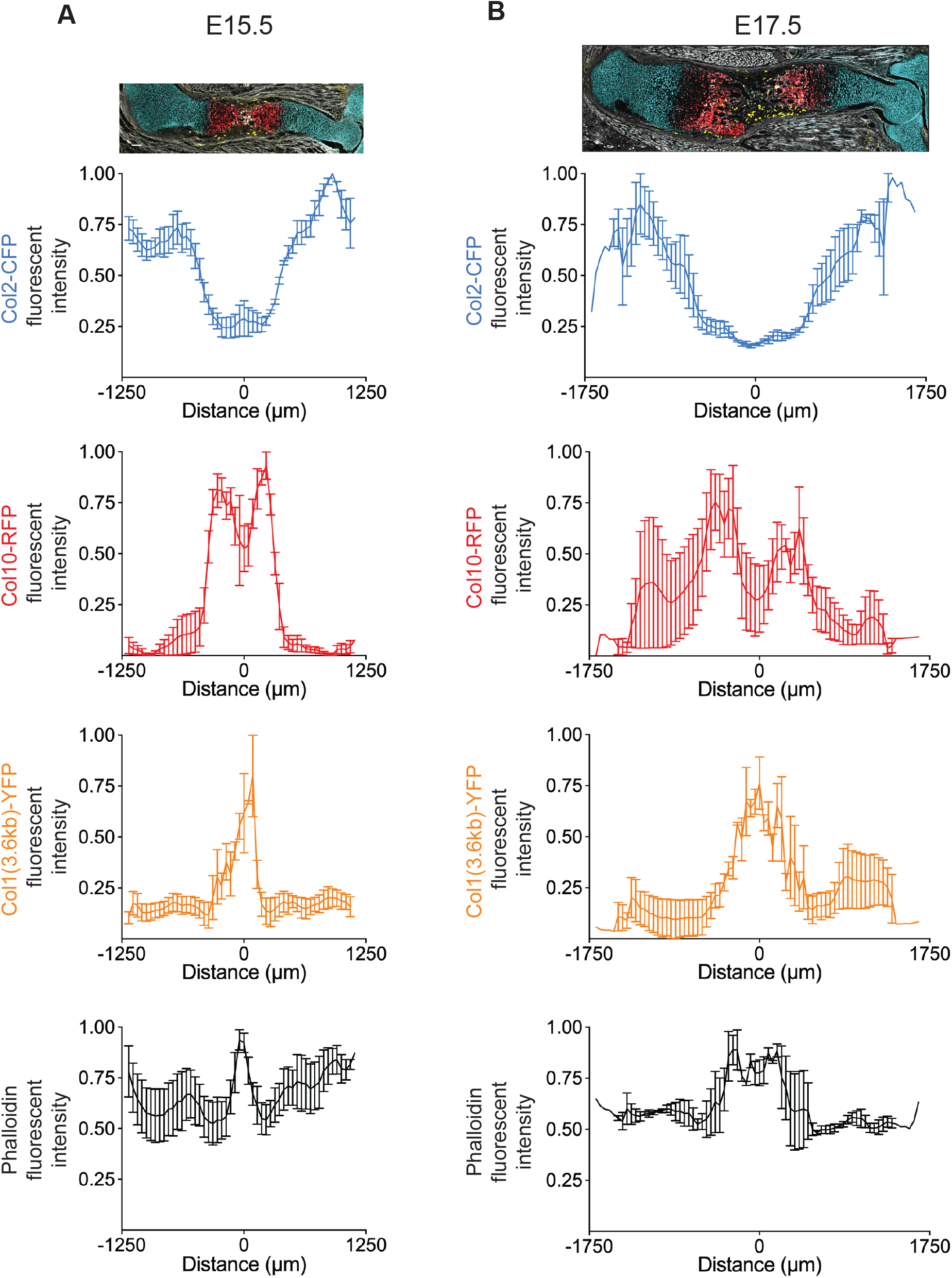
E15.5 and E17.5 Col2-CFP; ColX-RFP; Col1(3.6)-YFP intensity line plots by individual channel. (A) E15.5. (**B**) E17.5. Images are scaled to the plots. Images shown are repeated from figure 2 for clarity of the individual channel plots.

**Figure S3.**
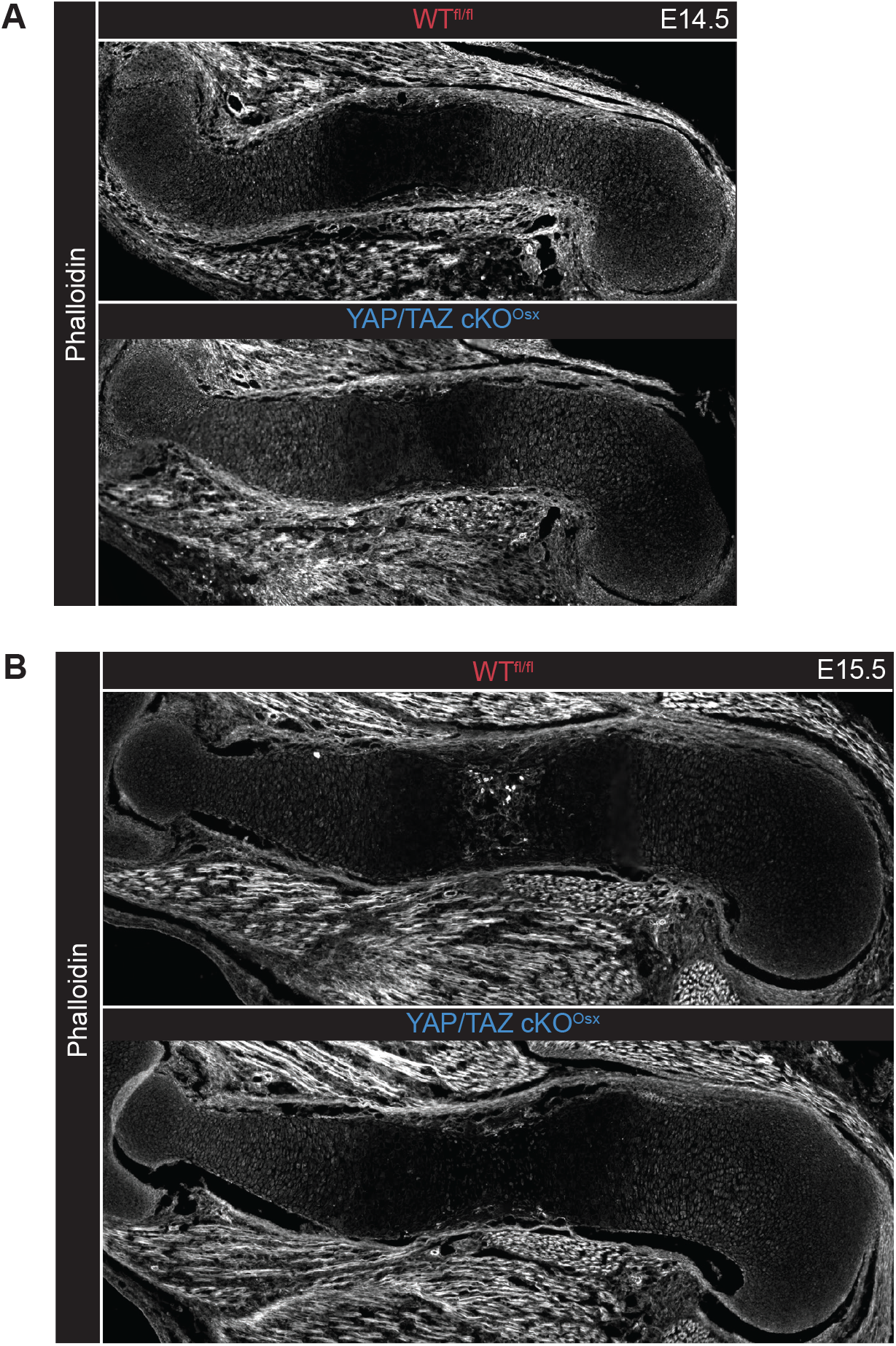
Phalloidin stained E14.5 and E15.5 WT^fl/fl^ and YAP/TAZ cKO^Osx^. (**A**) E14.5 humeri. (**B**) E15.5 humeri

**Figure S4.**
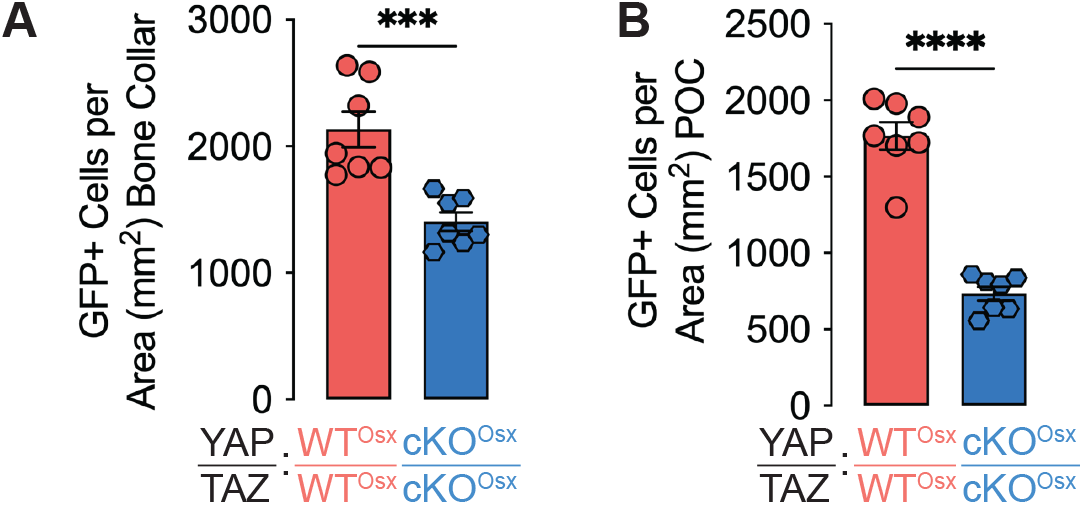
Osx::GFP^+^ cell density in E17.5 humeri. (**A**) Osx::GFP^+^ cell density in the bone collar. (**B**) Osx::GFP^+^ cell density in the primary ossficiation center (POC).

**Figure S5.**
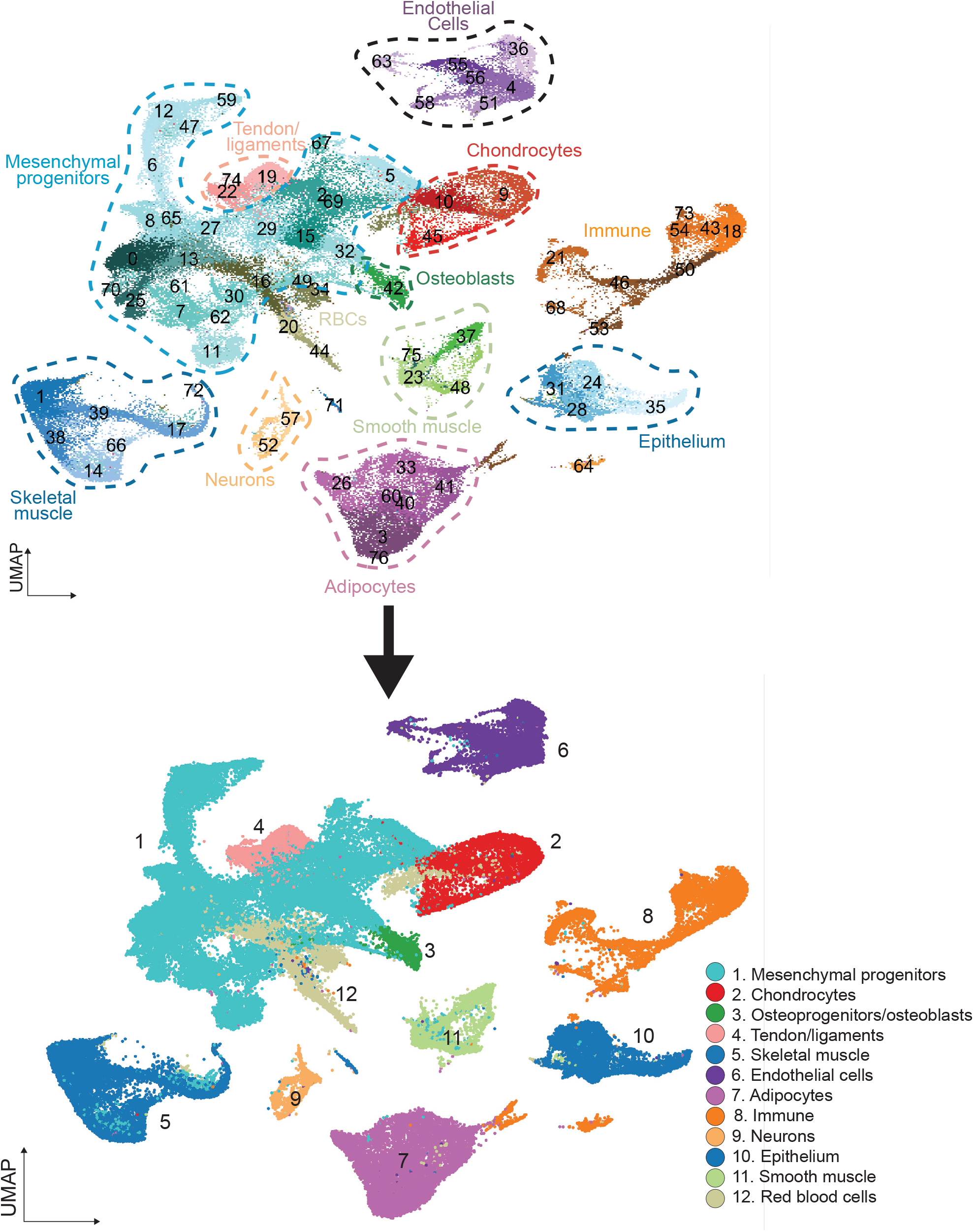
Cell clusters in the whole limb merged into major cell types. Louvain clustering identified 77 clusters in the whole fetal limb. We merged these clusters into 12 major cell types.

**Figure S6.**
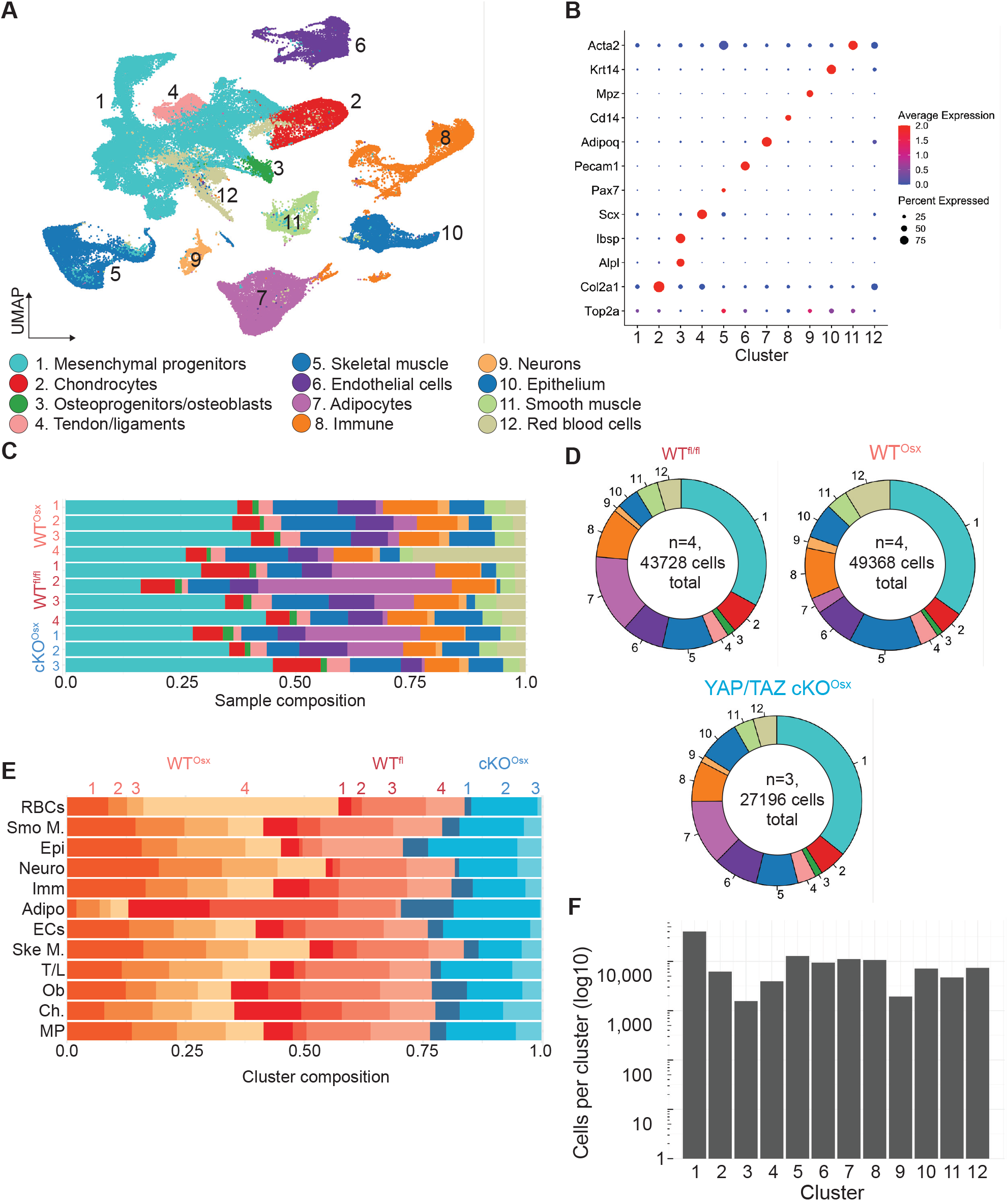
Major cell types in the fetal forelimb. (**A**) UMAP visualization showing cell types in from WT^fl/fl^, WT^Osx^, YAP/TAZ cKO^Osx^ fetal forelimbs. (**B**) Dotplot showing expression of canonical cell type markers. (**C**) Bar graph showing cell type proportions by sample. (**D**) Pie charts showing cell type proportions by genotype. The numbers in the middle of the pie chart indicates the genotype sample size and total high quality cells from each genotype analyzed here. (**E**) Bar graph showing sample proportions by cell type. (**F**) Bar graph showing the size of each major cell type cluster.

**Figure S7.**
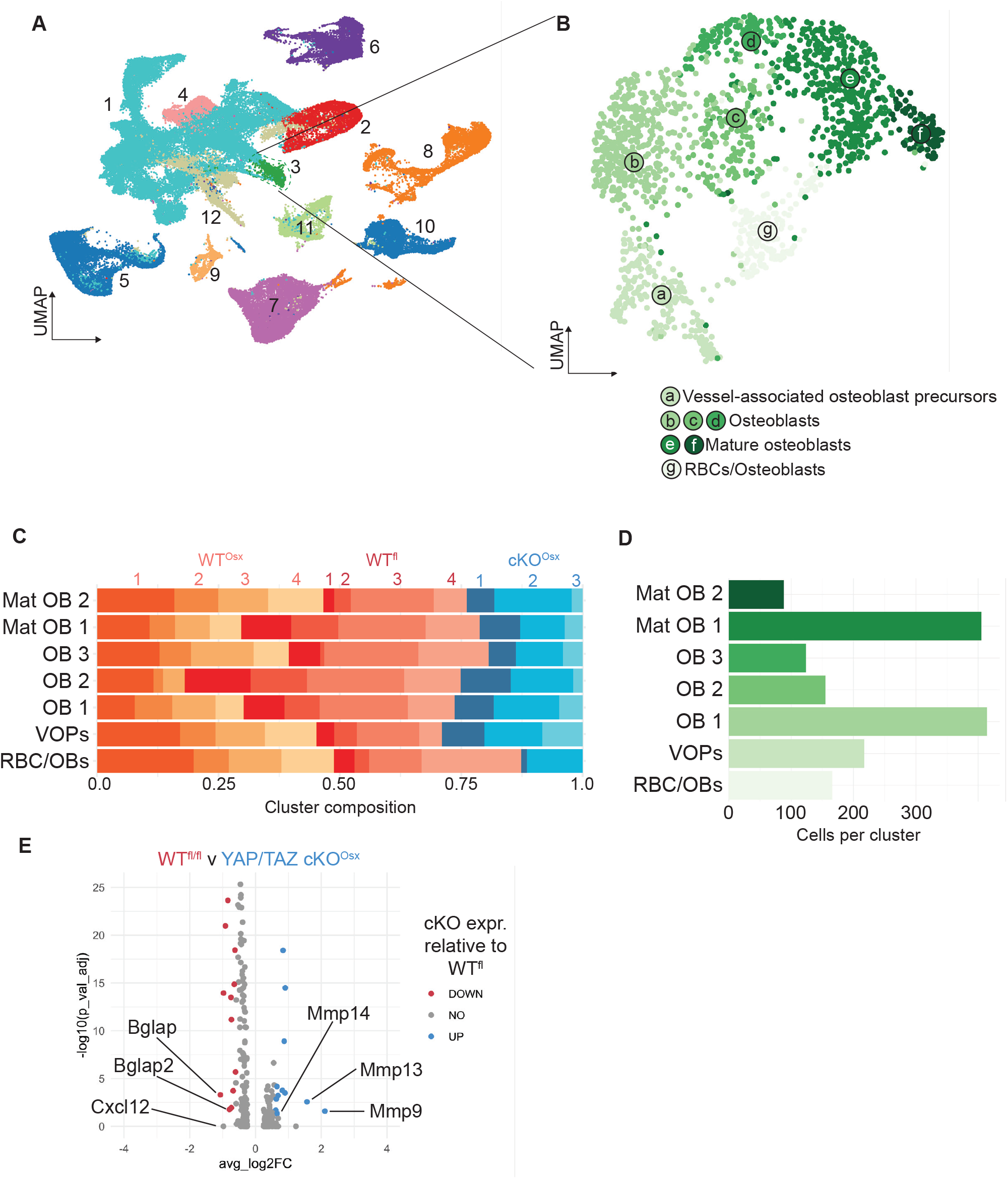
Osteoblast cell states. (**A**) UMAP visualization of fetal forelimb cells showing the osteoblasts highlighted. (**B**) UMAP visualization of osteoblast cell states. (**C**) Bar graph showing sample proportions by osteoblast cell state. (**D**) Bar graph showing the cluster size of each osteoblast cell state. (**E**) Volcano plot showing gene expression differences between WT^fl/fl^ and YAP/TAZ cKO^Osx^ osteoblasts overall. Red is decreased in the YAP/TAZ cKO^Osx^. Blue is increased in the YAP/TAZ cKO^Osx^.

**Figure S8.**
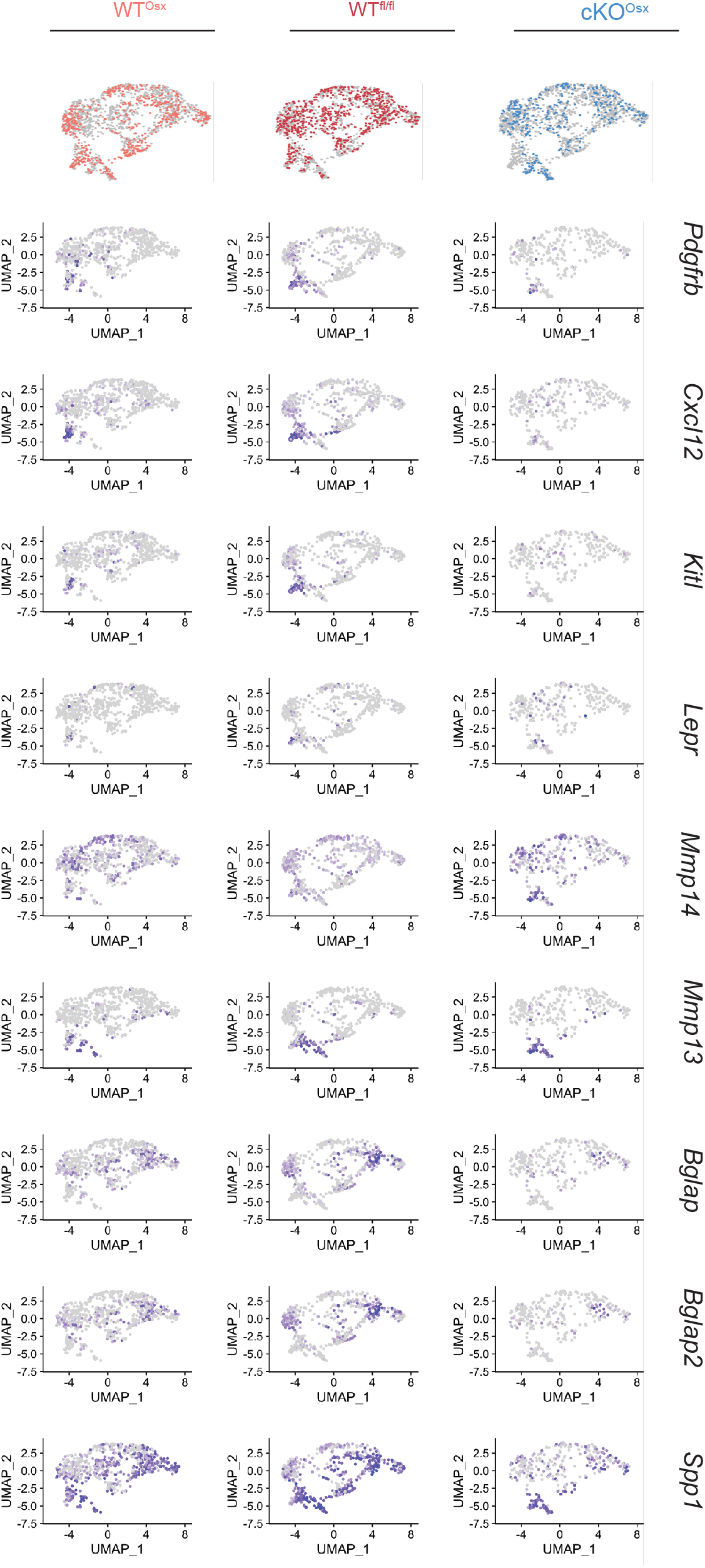
Gene plot visualization of gene expression in WT^Osx^, WT^fl/fl^, YAP/TAZ cKO^Osx^ in the osteoblast cell states. The top row shows the distribution of each genotype within the osteoblast cell states. The remaining rows show critically differentially expressed genes in the osteoblasts separated by genotype.

**Figure S9.**
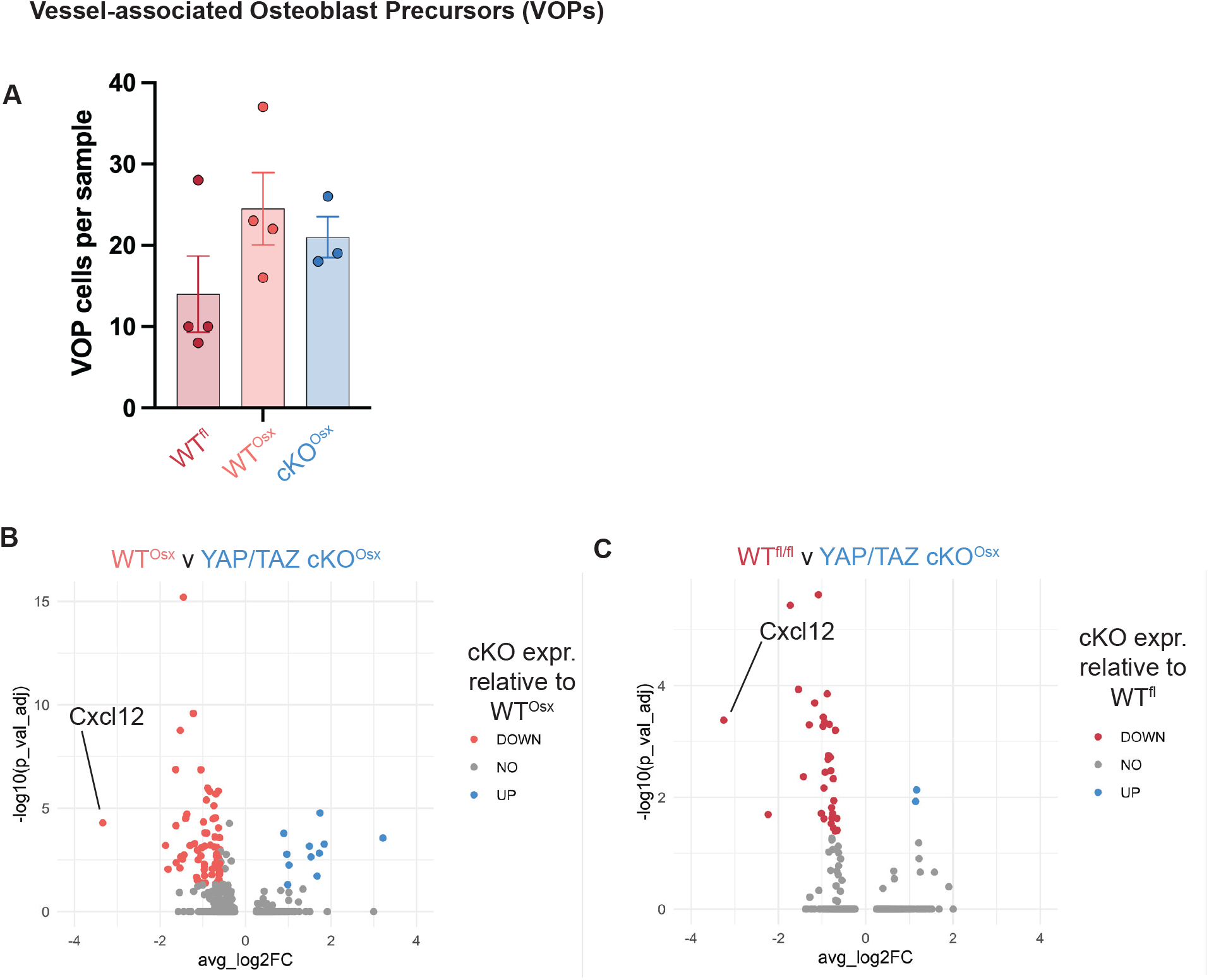
Supplemental data on the vessel associated osteoblast precursor (VOP) cell state. (**A**) Plot showing the number of VOPs in each sample, separated by genotype. (**B**) Volcano plot showing gene expression differences between WT^Osx^ and YAP/TAZ cKO^Osx^ VOPs. Red is decreased in the YAP/TAZ cKO^Osx^. Blue is increased in the YAP/TAZ cKO^Osx^. (**C**) Volcano plot showing gene expression differences between WT^fl/fl^ and YAP/TAZ cKO^Osx^ VOPs. Red is decreased in the YAP/TAZ cKO^Osx^. Blue is increased in the YAP/TAZ cKO^Osx^. *Cxcl12* is the most differentially reduced gene when YAP/TAZ is deleted from VOPs in comparison to both Wildtypes.

**Figure S10.**
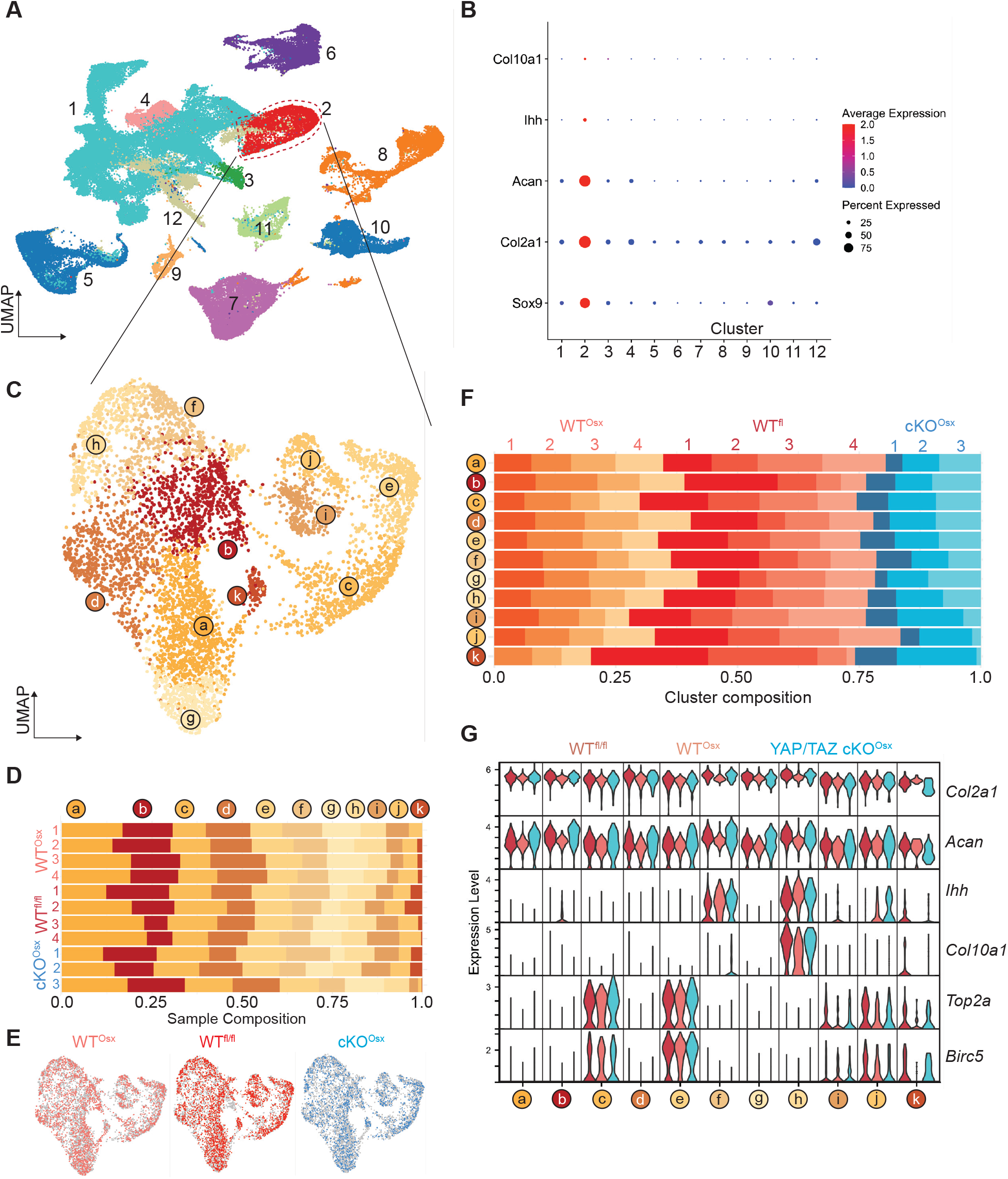
Chondrocyte cell states. (**A**) UMAP visualization of fetal forelimb cells showing the chondrocyte highlighted. (**B**) Dotplot showing chondrocyte marker gene expression among the whole fetal limb cell types **(C)** UMAP visualization of chondrocyte cell states. (**D**) Bar graph showing chondrocyte cell state proportions by sample. (**E**) UMAP visualization of chondrocyte cell states with each genotype highlighted. (**F**) Bar graph showing sample proportions by chondrocyte cell state. (**G**) Violin plot showing selected gene expression for each chondrocyte cell state by genotype.

**Figure S11.**
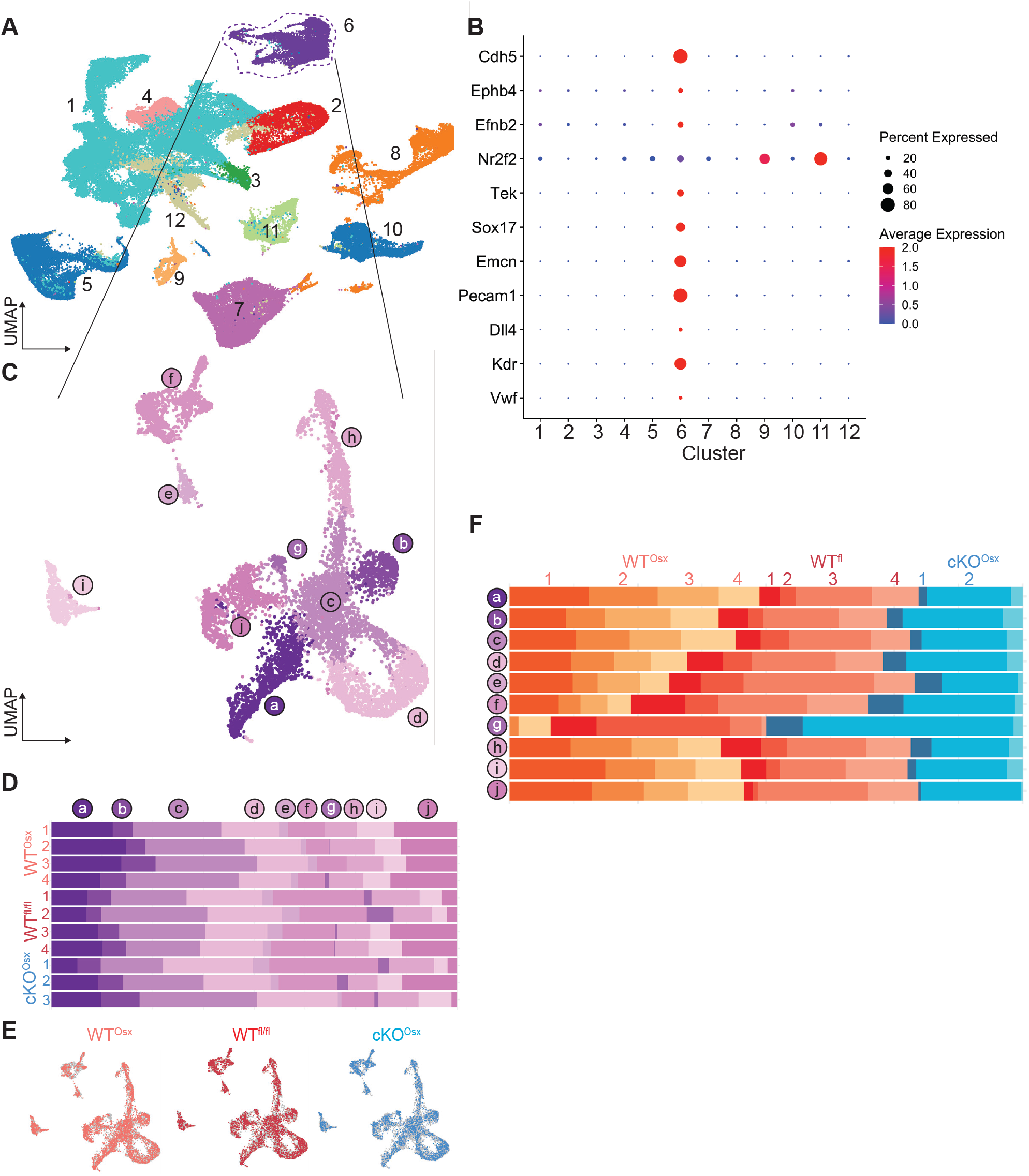
Endothelial cell states. (**A**) UMAP visualization of fetal forelimb cells showing the endothelial cells highlighted. (**B**) Dotplot showing endothelial marker gene expression among the whole fetal limb cell types (**C**) UMAP visualization of endothelial cell states. (**D**) Bar graph showing endothelial cell state proportions by sample. (**E**) UMAP visualization of endothelial cell states with each genotype highlighted. (**F**) Bar graph showing sample proportions by endothelial cell state.

**Figure S12.**
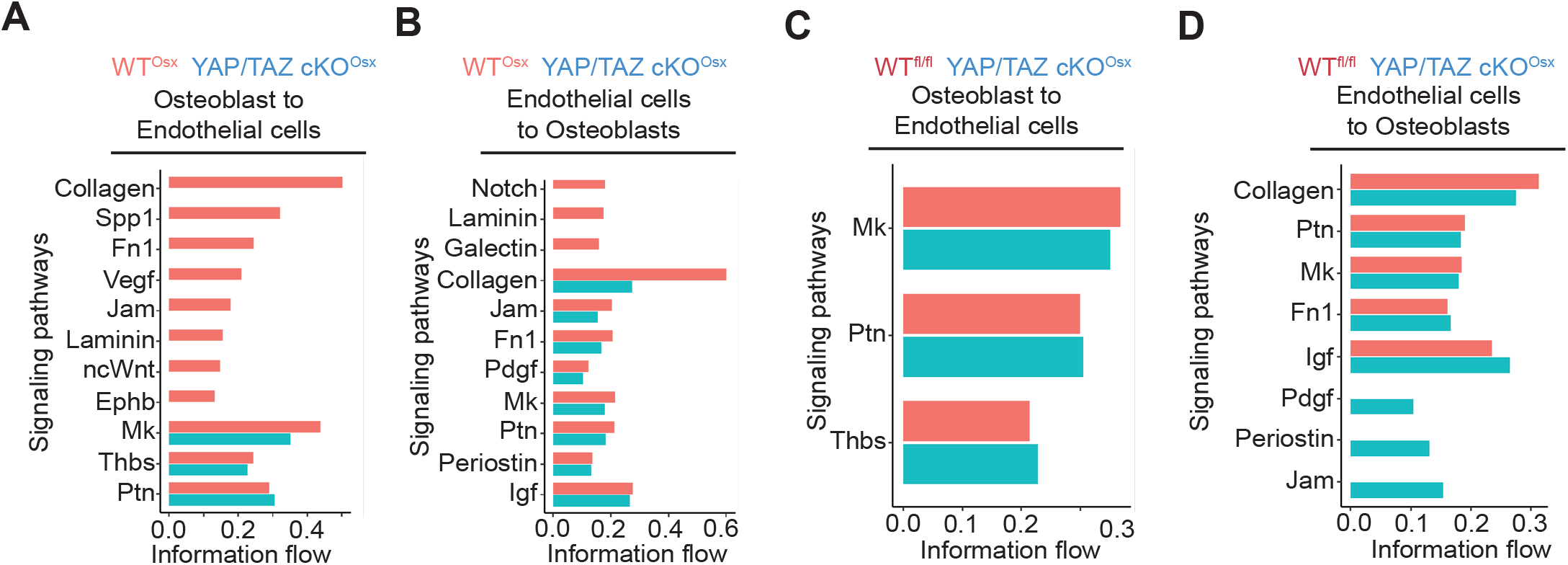
Cell-cell communication between osteoblast cell type and endothelial cell type by CellChat. (**A**) Cell-cell communication pathways from osteoblasts to endothelial cells in WT^Osx^ and YAP/TAZ cKO^Osx^. (**B**) Cell-cell communication pathways from endothelial cells to osteoblasts in WT^Osx^ and YAP/TAZ cKO^Osx.^ (**C**) Cell-cell communication pathways from osteoblasts to endothelial cells in WT^fl/fl^ and YAP/TAZ cKO^Osx^. (**D**) Cell-cell communication pathways from endothelial cells to osteoblasts in WT^fl/fl^ and YAP/TAZ cKO^Osx^.

**Figure S13.**
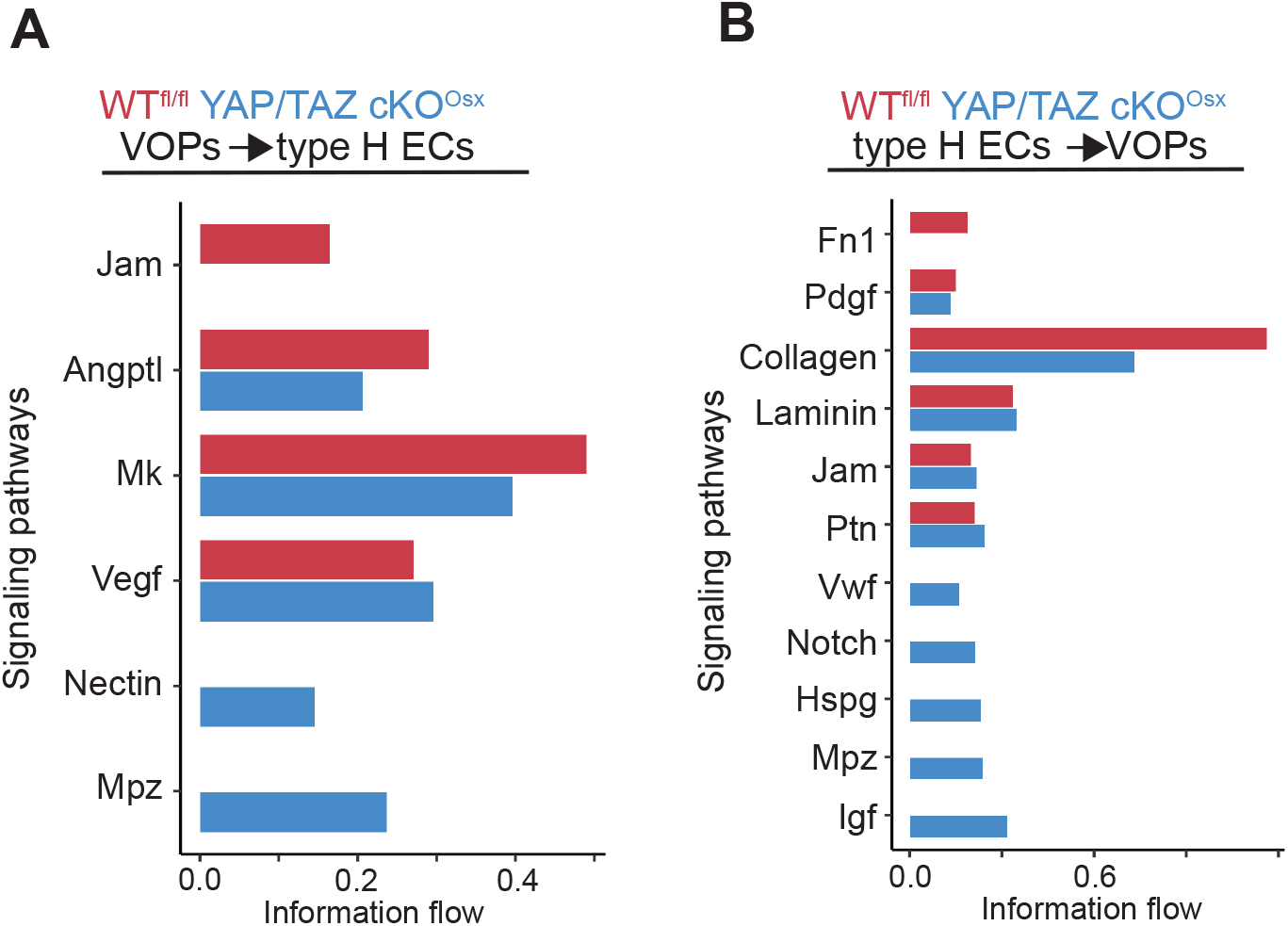
Cell-cell communication between Vessel associated osteoblast precursors (VOPs) and type H endothelial cells by CellChat. (**A**) Cell-cell communication pathways from VOPs to type H endothelial cells in WT^fl/fl^ and YAP/TAZ cKO^Osx^. (**B**) Cell-cell communication pathways from type H endothelial cells to VOPs in WT^fl/fl^ and YAP/TAZ cKO^Osx^.

**Figure S14.**
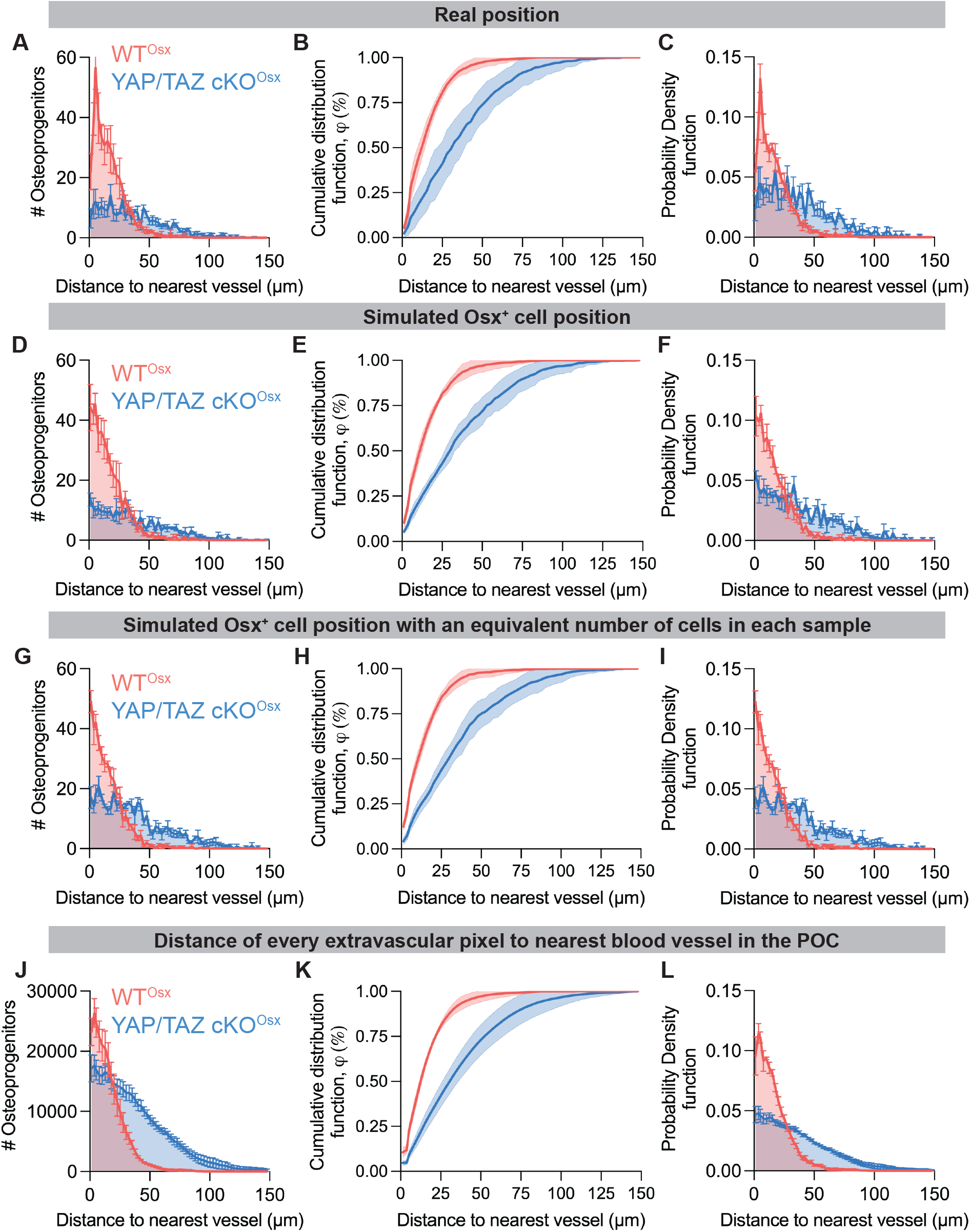
Osx::GFP+ cell proximity to Endomucin^+^ blood vessels and computational varied tests. (**A-C**) (A) histogram, (B) Cumulative distribution function, (C) Probability density function for the real distances from each Osx::GFP+ cell to its nearest blood vessel in the primary ossification center of WT^Osx^ and YAP/TAZ cKO^Osx^. (**D-F**) (D) histogram, (E) Cumulative distribution function, (F) Probability density function for the first computationally varied condition, in which the position of each Osx::GFP+ cell is randomized within the extravascular space of its respective primary ossification center. Randomizing the position does not collapse the difference in proximity distribution between WT^Osx^ and YAP/TAZ cKO^Osx^ samples, thus Osx::GFP+ cell position doesn’t explain the differences. (**G-I**) (G) histogram, (H) Cumulative distribution function, (I) probability density function for the second computationally varied condition, in which a fixed number (400) of Osx::GFP+ cells are positionally randomized within the extravascular space of its respective primary ossification center. Fixing the number of Osx::GFP+ cells and randomizing their position does not collapse the difference in proximity distribution between WT^Osx^ and YAP/TAZ cKO^Osx^ samples, thus Osx::GFP+ cell density doesn’t explain the differences. (**J-L**) (J) histogram, (K) Cumulative distribution function, (L) probability density function for the third computationally varied condition, in which the distance of every extravascular pixel to its nearest vessel was calculated.

**Figure S15.**
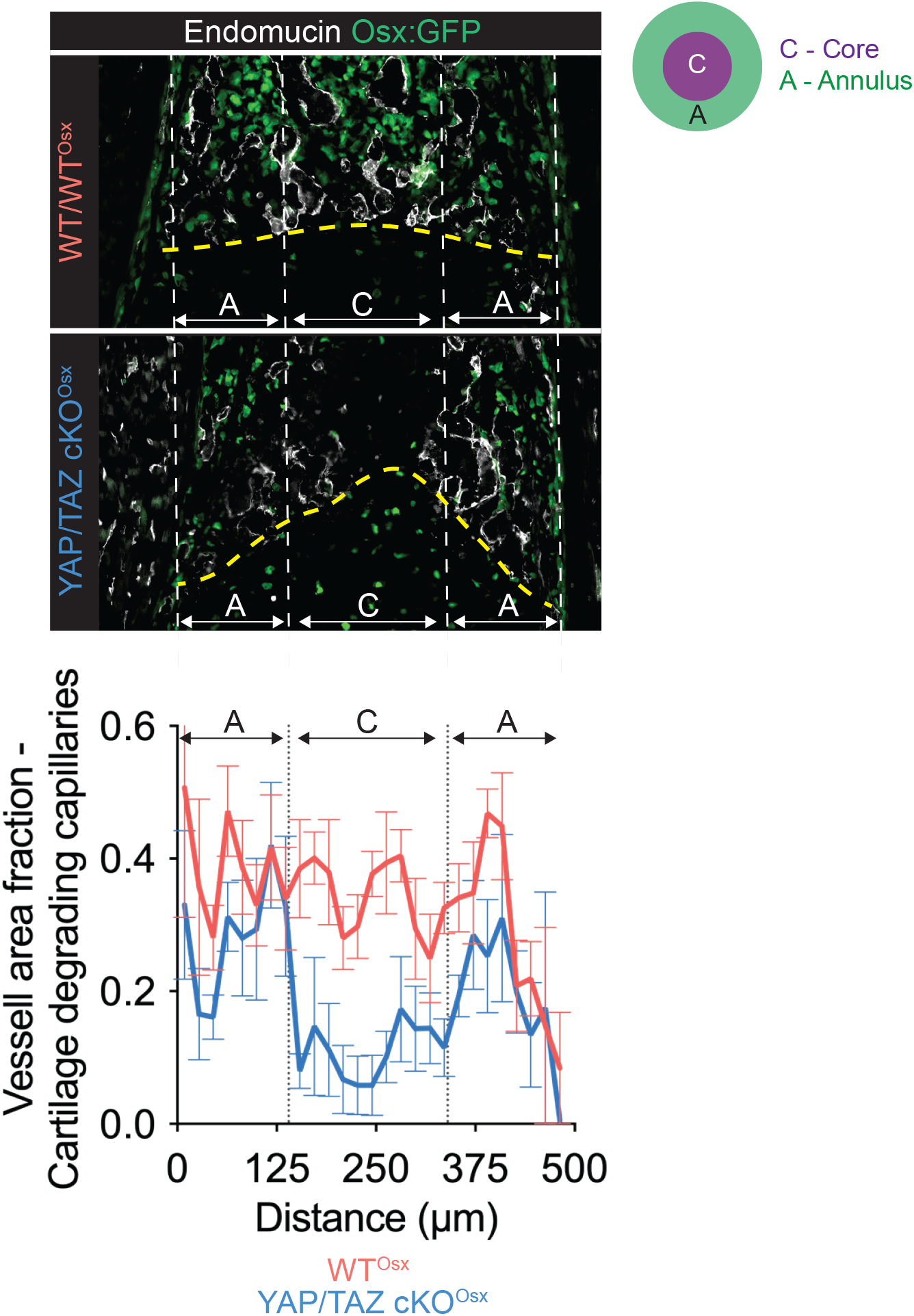
Quantification of the blood vessels within 50μm of the chondro-osseous junction in the core and outer annulus. C – Core. A -Annul

**Figure S16.**
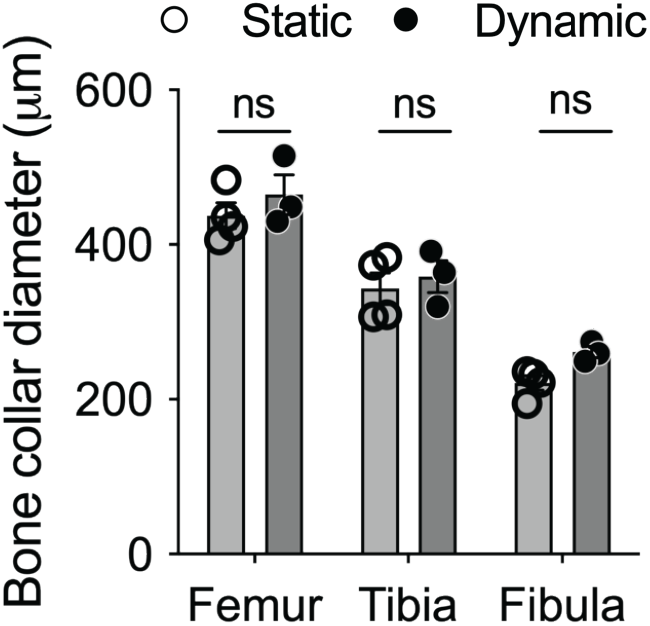
Bone collar diameter of explant C57Bl6 limbs in the cultured in the mechanostimulation bioreactor.

**Figure S17.**
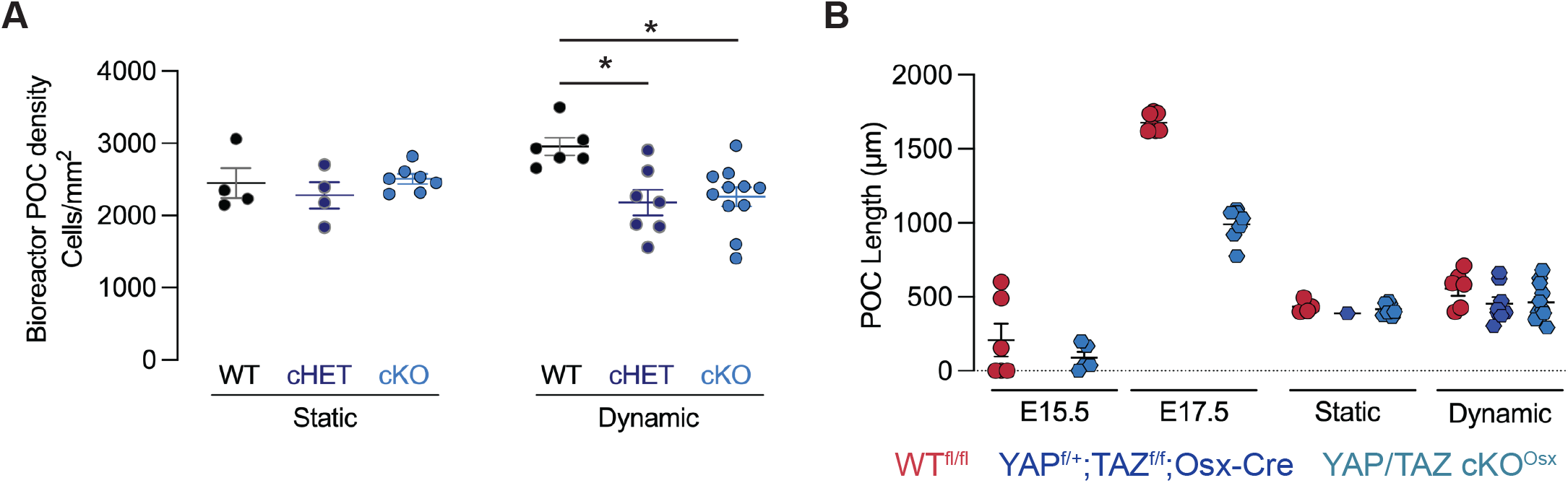
Supplemental data for the genetic Osx-conditional YAP/TAZ deletion bioreactor experiments. **(A)** Primary ossification center (POC) cell density after 6 days of culture in the mechanostimulation bioreactor. **(B)** POC length in development at E15.5 and E17.5 and in E15.5+6d culture under static or dynamic conditions.

